# Surprising variation in the outcome of two malaria genetic crosses using humanized mice: implications for genetic mapping and malaria biology

**DOI:** 10.1101/2019.12.13.871830

**Authors:** Katrina A. Button-Simons, Sudhir Kumar, Nelly Carmago, Meseret T. Haile, Catherine Jett, Lisa A. Checkley, Spencer Y. Kennedy, Richard S. Pinapati, Douglas A. Shou, Marina McDew-White, Xue Li, François H. Nosten, Stefan H. Kappe, Timothy J. C. Anderson, Jeanne Romero-Severson, Michael T. Ferdig, Scott J. Emrich, Ashley M. Vaughan, Ian H. Cheeseman

## Abstract

Genetic crosses are most powerful for linkage analysis when progeny numbers are high, when parental alleles segregate evenly and, for hermaphroditic organisms, when numbers of inbred progeny are minimized. We previously developed a novel genetic crossing platform for the human malaria parasite *Plasmodium falciparum*, an obligately sexual, hermaphroditic protozoan, using mice carrying human hepatocytes (the human liver-chimeric FRG NOD huHep mouse) as the vertebrate host. Here we examine the statistical power of two different genetic crosses – (1) between a laboratory parasite (NF54) of African origin and a patient-derived Asian parasite, and (2) between two sympatric patient-derived Asian parasites. We generated >140 unique recombinant clones over a 12-month period from the four parental genotypes, doubling the number of unique recombinant progeny generated in the previous 30 years. Both crosses show bi-parental inheritance of plastid markers amongst recombinant progeny, in contrast to previous crosses (conducted using chimpanzee hosts) which carried single dominant plastid genotypes. Both crosses show distinctive segregation patterns. The allopatric African/Asian cross has minimal levels of inbreeding (2% of clonal progeny are inbred) and extreme skews in marker segregation, while in the sympatric Asian cross, inbred progeny predominate (66% of clonal progeny are inbred) and parental alleles segregate evenly. Using simulations, we demonstrate that these progeny arrays (particularly the sympatric Asian cross) have excellent power to map large-effect mutations to a 31 kb interval and can capture complex, epistatic interactions that were far beyond the capacity of previous malaria crosses to detect. The extreme segregation distortion in the allopatric African/Asian cross erodes power to detect linkage in several genome regions, but the repeatable distortions observed offer promising alternative approaches to identifying genes underlying traits of interest. These crosses show surprising variation in marker segregation, nevertheless, the increased progeny numbers improve our ability to rapidly map biomedically important parasite traits.

**Author Summary:** Understanding how genome mutations contribute to newly emerging drug resistance in parasites like *Plasmodium falciparum* is important to monitor the spread of drug resistance. This scenario has been playing out in Southeast Asia with the emergence and spread of artemisinin resistance. Here we show that new *P. falciparum* genetic crosses, using mice carrying human liver cells and infused with human red blood cells (the human liver-chimeric FRG NOD huHep/huRBC mouse), provide an important new tool for understanding complex interactions underlying drug resistance phenotypes. We report two new genetic maps with 84 and 60 unique recombinant progeny, which doubles the number of progeny available from 4 previous *P. falciparum* genetic crosses. Through extensive simulations we show that with 84 progeny we can find association for a gene that controls only 20% of the variation in a phenotype. We also show that a cross generated from Southeast Asian parasites collected from the same geographic region have unique characteristics not previously observed in *P. falciparum* genetic crosses. This Southeast Asian cross exhibits even segregation across the genome, unbiased inheritance of mitochondria and apicoplast and higher levels of inbreeding than previously observed.

## Introduction

Eukaryotic parasites inflict a high burden of morbidity and mortality particularly in the developing world. Control of these pathogens is threatened by drug resistance [1, 2]. Understanding the genetic architecture of drug resistance in eukaryotic pathogens is essential to understand treatment failure. Previous studies in Plasmodium, Trypanosome and Leishmania parasites revealed the genetic architecture of drug resistance is unexpectedly complex [3–6]. For example, emergent artemisinin resistance in the human malaria parasite, *Plasmodium falciparum,* has been causally associated with multiple independent mutations in one gene, *pfK13,* which explain nearly all the variation in this phenotype [7–9]. However, mutations in the *pffd*, *pfarps10*, *pfmdr2*, and *pfcrt* genes are significantly associated with resistance, and have been proposed to constitute a genetic background highly predisposed to the development of resistance [7]. Several techniques have been used to identify the genetic determinants of complex phenotypes in eukaryotic pathogens including GWAS [7, 10], *in vitro* selections [8], QTL analysis in controlled genetic crosses [11–14] and bulk segregant analysis [5, 15]. Controlled genetic crosses offer a uniquely powerful way to dissect the genetic architecture of a complex trait. For example, the F_1_ progeny of a controlled cross revealed that *P. falciparum* sensitivity to quinine was associated with loci on chromosomes 5, 7 and 13, with the chromosome 5 and 7 loci containing known drug resistance transporters *pfcrt and pfmdr1* [3].

*P. falciparum* has the potential to be a particularly powerful genetic mapping system because of its unusually high recombination rate of 11-13.3 kb/cM [13, 16, 17], a haploid state for most of the life cycle, and the ability to clone every F_1_ progeny *in vitro*, creating effectively immortal mapping populations in a single generation. Also, *P. falciparum* has a small genome (23 Mb) and a high-quality reference assembly [18] with frequent annotation updates [19, 20]; consequently, re-sequencing and comprehensive analysis of the genome of F_1_ progeny is simple and cost effective [21]. Generating controlled genetic crosses in *P. falciparum*, however, has historically been a difficult and time-consuming process requiring splenectomized chimpanzees in place of a human host. This has resulted in only four genetic crosses being performed over a thirty-year period. F_1_ mapping populations from all four previous *P. falciparum* genetic crosses have been small, containing 33, 35, 15 [21] and most recently 27 individual recombinant progeny [13]. When compared to the thousands of F_1_ progeny possible in many plants and fruit flies [22], these numbers are small indeed. To use genetic mapping to elucidate the genetic architecture of emerging drug resistance in *P. falciparum* we need to be able to rapidly create genetic crosses with large numbers of progeny from recent field isolated parasites which exemplify highly relevant clinical traits such as drug resistance.

Here we report the production of large numbers of unique recombinant progeny from human liver-chimeric FRG huHep mice infused with human red blood cells. Although these mice were previously reported as an option for producing new *P. falciparum* genetic crosses once chimpanzee research was discontinued [23], until now they have failed to produce more progeny than historic crosses. In this paper, we successfully produced two new genetic crosses in under twelve months using recent clinically derived *P. falciparum* isolates with emerging resistance phenotypes. This effort was aided by a new progeny characterization bioinformatics framework that filters SNP variants and identifies clonal unique recombinant progeny. We generate genetic maps for each cross (84 and 60 unique recombinant progeny, respectively) and provide the most detailed investigation of inbreeding, plastid inheritance, and cross-over rates in malaria parasite genetics to date. One cross exhibited abundant segregation distortion. We confirm this is repeatable by independently replicating the genetic cross, and exclude a fluorescent marker integrated into genome of one parent as the cause. Through simulation and mapping with real data we investigate the power to detect genetic associations as a function of the number of progeny. We also examine the effect of segregation distortion on power in mapping a phenotype in a cross with varying levels of segregation distortion.

## Results

### Rapid Generation of Genetic Crosses

Over a 12-month period we carried out two independent genetic crosses. The first between a laboratory-adapted African line (NF54) and a newly cloned clinical isolate (NHP4026) from the Thai-Myanmar border, the second between two newly cloned clinical isolates (MKK2835 and NHP1337) from the Thai-Myanmar border. These crosses yielded 84 and 60 clonal unique recombinant progeny lines respectively. The pipeline to the point of analyzing recombinant progeny is technically challenging and takes approximately six months (Fig 1). Initially, we confirmed that the clinical isolate parental lines produced infectious gametocytes that gave rise to infectious sporozoites that could the successfully infect the liver of human hepatocyte-chimeric FRG NOD huHep mice and subsequently transition to *in vivo* and then *in vitro* blood stage culture. After this confirmation, the steps to successfully complete a genetic cross includes asexual culture and expansion, gametocyte maturation, mixing of parental gametocytes and transmission to mosquitoes, confirmation of successful mosquito stage development, salivary gland sporozoite isolation and infection of human hepatocytes in the FRG NOD huHep mouse, liver stage development, infusion of human red blood cells, the *in vivo* transition from liver stage-to-blood stage, the subsequent transition to *in vitro* blood stage culture coupled with cloning by limiting dilution and finally clonal expansion, confirmation of clonality and genome sequencing of recombinant progeny (Fig 1).

**Fig 1.**
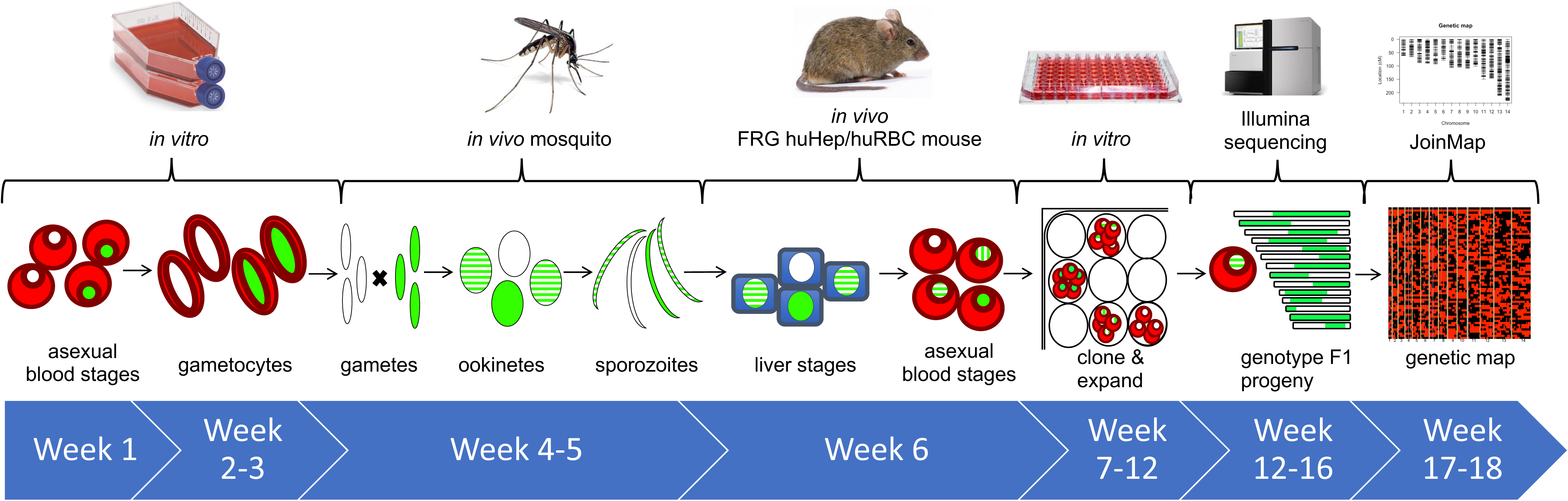
Timeline for performing *P. falciparum* crosses in FRG huHep/huRBC mice. Uncloned F1 progeny from *P. falciparum* genetic crosses of recent field isolates can be recovered in 6 weeks from asexual stage culture of parent lines. Cloning of potential F1 recombinant progeny takes an additional 6 weeks. Next generation sequencing of potential recombinant progeny and identification of unique recombinants via our new pipeline takes an additional 6 weeks.

In total we initiated three independent crosses using five parental genotypes (NF54- GFPLuc x NHP4026, NF54WT x NHP4026 and MKK2835 x NHP1337). The second of these crosses was performed to test if a GFP cassette integrated into the genome had driven a peak in segregation distortion (described below). The progeny from the first crosses were combined (subsequently referred to as NF54 x NHP4026) to form one genetic map (described below). We set up each genetic cross by infecting multiple cages of mosquitos with mixed gametocyte cultures of our parental lines (S1 Table). Details NF54-GFPLuc x NHP4026 were previously published [23]. For NF54WT x NHP4026 three cages were used to infect three mice by IV injection or mosquito bite (MB) (one cage per mouse). Two mice were infected by MB using cages with 250 mosquitos with prevalence of 73% and 58% and median 3 oocyst/mosquito. One mouse was infected by IV injection of 1 million sporozoites dissected from 250 mosquitos with infection prevalence of 73% and median 6 oocysts/mosquito. Assuming no attrition in parasite genotypes, and a perfect outcrossing rate this would mean that 2190 and 1740 unique recombinant progeny respectively were possibly inoculated into two mice by mosquito bite and a pool of 1460 unique recombinant progeny was used to infect one mouse via IV infection. Similarly, for MKK2835 x NHP1337 four cages of mosquitos were infected with pools of MKK2835 and NHP1337 gametocyes and the cage with the best infections (80% prevalence and median 3 oocysts/mosquito) was used to infect a single mouse via IV injection with 2.7 million sporozoites. We would expect a maximum of 1958 unique recombinant progeny based upon 80% successful infections and a median of 3 oocysts per mosquito and 204 mosquitos.

### Numbers of Unique Recombinant Progeny

In these malaria parasite crosses, the F_1_ progeny are present in the blood of the infected FRG NOD huHep/huRBC mouse and must be isolated by limiting dilution cloning. The progeny isolated in this way are not guaranteed to be clonal because a small subset of post-dilution cultures will have been initiated with more than one clone. Additionally, as the malaria parasite undergoes clonal expansion in the mosquito, liver and mouse blood stream [24] we may sample the same recombinant genotype more than once. Since the parents in both crosses readily produce fertile male and female gametocytes it is also possible for selfed progeny to be produced. We thus developed a bioinformatics pipeline to identify clonal unique recombinant F_1_ progeny filtering out non-clonal progeny, selfed progeny and repeat sampling of the same genotype (see Methods).

Genetic characterization of previous crosses was initially carried out with RFLP or MS markers [16, 25] and unique recombinant progeny from these crosses were recently sequenced to create a community resource [21]. For NF54 x NHP4026, we filtered out some non-unique recombinant progeny using MS genotyping and then performed direct genome sequencing of cloned parasites. For MKK2835 x NHP1337, we performed genome sequencing of all cloned parasites. For each prospective progeny, sequencing reads were mapped to version 3 of the *P. falciparum* genome [26] and SNP variants were called jointly across parents and perspective progeny and filtered to contain SNPs in the 20.8 Mb core genome as defined in Miles et al. 2016 [21].

In NF54 x NHP4026, 10,472 high-quality bi-allelic SNPs (1 SNP per 2.0 kb) differentiate the two parents. For this cross, 166 prospective progeny were identified during limiting dilution cloning. After filtering to remove non-clonal and selfed progeny 128 recombinant progeny remained (Fig 2), 84 of which were unique. In MKK2835 x NHP1337, the parent lines are sympatric patient-derived Asian parasites. Despite their higher degree of relatedness we identified 7,198 high-quality bi-allelic SNPs (1 SNP per 2.9 kb) that distinguish the two parents. For this cross 266 prospective progeny were identified during limiting dilution cloning. Filtering was performed to remove non-clonal and selfed progeny leaving 61 recombinant progeny (Fig 2), 60 of which were unique. We initiated multiple cloning rounds to maximize the capture of unique recombinant progeny from each cross. Interestingly, across all crosses each cloning round produced nearly distinct sets of recombinant progeny, with only one repeat genotype across cloning rounds (Fig 2 and S1 Fig).

**Fig 2.**
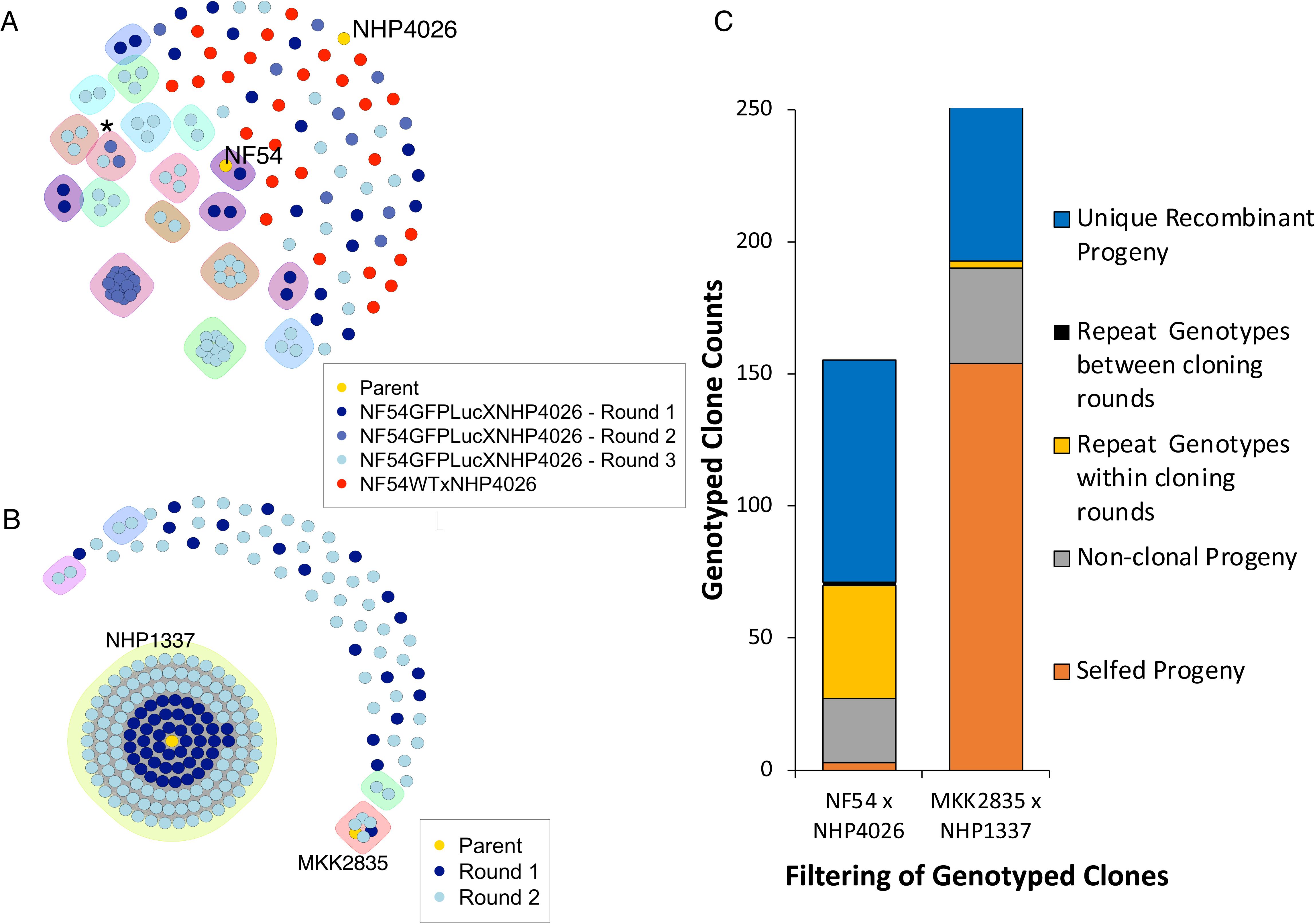
Cloning results and estimated recombinant progeny for each cross. (A and B) Genotyping results for each cross, clusters denote individual clones of the same genotype. (A) NF54 x NHP4026 contained few selfed progeny and almost all repeat sampling of the same genotype (clusters) occurred with a cloning round, * denotes the only observed repeat sampling event between cloning rounds. (B) MKK2835 x NHP1337 produced many selfed progeny and few instances of repeat sampling of recombinant genotypes all from cloning round 2. (C) Progeny for NF54 x NHP4026 and MKK2835 x NHP1337 cross were filtered to identify unique recombinant progeny (blue). Selfed progeny (orange), non-clonal progeny (grey) and repeat sampling of the same genotype within a cloning round (yellow) and between cloning rounds (black) were filtered out of total genotyped progeny.

### Inbreeding, Outbreeding and Plastid inheritance

Through our filtering process we identified stark differences in patterns of outcrossing between these two crosses. The clones recovered from NF54 x NHP4026 contained few selfed progeny with three selfed NF54 progeny and 0 selfed NHP4026 progeny (1.8%, 3/166 progeny selfed). In contrast, in MKK2835 x NHP1337 we observed a large amount of selfing with 144 selfed NHP1337 progeny and five MKK2835 selfed progeny (56%, 149/266 progeny selfed; Fig 2). In both crosses, when cloning was initiated immediately after mouse exsanguination or within five days of establishing *in vitro* culture, almost all recombinants were unique (S2 Table and S1 Fig). Interestingly, when cloning was initiated within five days, whether from continuous *in vitro* culture or cryopreservation of bulk culture, the percentage of recombinants that were unique was high (90-100% for continuous culture vs. 93% from a thawed cryopreserved bulk culture). However, when cloning was initiated after 14 or 19 days of *in vitro* culture from cryopreserved bulk culture, a lower percentage of unique recombinant progeny were recovered with 46% and 50% of recombinants identified as unique (S2 Table and S1 Fig).

*P. falciparum* parasites contain two plastid genomes, the mitochondria and apicoplast, both of which are maternally inherited [27]. Despite *P. falciparum* being hermaphroditic, in previous genetic crosses nearly all plastid genomes in the progeny originated from a single parent [28, 29]. We show here that this is not the case. In each cross we observed both plastid genotypes among the unique recombinant progeny. After excluding selfed genotypes we observed 17.9% NF54 plastid genotypes in NF54 x NHP4026 and 41.7% MKK2835 plastid genotypes in MKK2835 x NHP1337.

### Genetic maps and recombination rates

For each genetic cross, we generated a genetic map (S3 and S4 Tables) using JoinMap v4.1 from phased genotype data for all unique recombinant progeny (see Methods). The map size for both crosses is consistent with map lengths reported for previous crosses (1521 cM for NF54 x NHP4026 and 1626 cM for MKK2835 x NHP1337, Table 1). The recombination rate was 13.7 kb/cM for NF54 x NHP4026 and 12.8 kb/cM for MKK2835 x NHP1337, which were comparable to the range observed in previous crosses (Table 1). In NF54 x NHP4026 genetic map, markers initially sorted into 13 linkage groups, with each representing markers known to reside on single chromosomes, with the exception of one linkage group which contained all markers on chromosome 7 and 14. Adjusting joinMap parameters resulted in separating the 13^th^ linkage group into 2, recovering distinct sets for chromosomes 7 and 14. In MKK2835 x NHP1337 all markers coalesced into 14 linkage groups which exactly corresponded to chromosomes.

**Table 1.**

| Cross | F<sub>1</sub> Progeny Number | Genetic Map Length | Recombination Rate |
| --- | --- | --- | --- |
| HB3 x Dd2 | 35<sup>[16]</sup> | 1556<sup>[16]</sup> | 12.1 kb/cM<sup>[16]</sup> |
| 3D7 x HB3 | 15<sup>[21]</sup> |  | 11 kb/cM<sup>[30]</sup> |
| 7G8 x GB4 | 32<sup>[17]</sup> | 1655<sup>[17]</sup> | 12.8 kb/cM<sup>[17]</sup> |
| GB4 x 803 | 27<sup>[13]</sup> |  | 13.3 kb/cM |
| NF54 x NHP4026 | 84 | 1521 | 13.7 kb/cM |
| MKK2835 x NHP1337 | 60 | 1626 | 12.8 kb/cM |

To generate a graphic display of the physical map, 5 kb windows of the core genome were phased to indicate inheritance blocks for each unique recombinant progeny (Fig 3A and 3B). NF54 x NHP4026 shows sections of the genome where inheritance is dominated by one parental genotype or the other (Fig 3A). In contrast the physical recombination map for MKK2835 x NHP1337 shows a more even inheritance pattern across the genome (Fig 3B).

**Fig 3.**
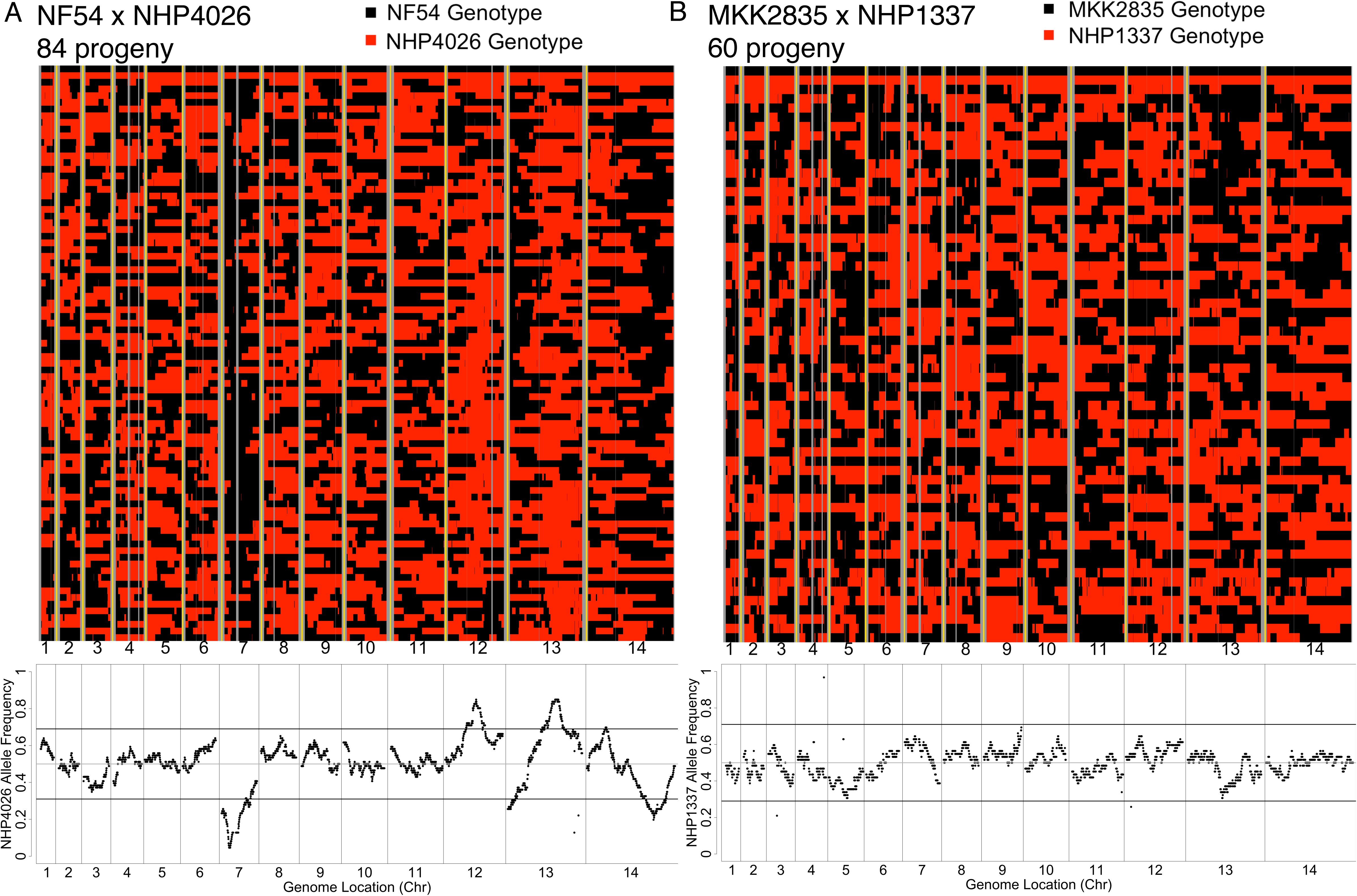
Physical maps for crosses. Physical maps (A & B) depict inheritance patterns in 5KB blocks for each progeny (y axis) across core regions of the 14 nuclear chromosomes (x axis) with black representing the drug susceptible parent and red the drug resistant parent, non-core regions of the genome with no variant calls are shown in grey with yellow showing chromosome boundaries. (A) The physical map for NF54 x NHP4026 shows several regions where haplotype blocks are primarily inherited from either parent and deviate significantly from the expected 1:1 ratio. (B) The physical map for MKK2835 x NHP1337 shows more even inheritance ratios across the genome with no significant deviations from expected mendelian ratios.

### Repeatability of segregation distortion

We observe regions with significant segregation distortion (chi squared test for deviation from expected Mendelian ratio of 1:1, p<0.001) in NF54 x NHP4026 that are consistent in both replicates (Fig 3A and 4A). In contrast, we observe no significant segregation distortion in MKK2835 x NHP1337 (Fig 3B). Specifically, in both replicates of NF54 x NHP4026 we observe replicated significant deviations from the Mendelian expectation of 1:1 inheritance on chromosomes 7, 12, 13 and 14 (Fig 3A) with a concordance correlation coefficient of 0.66 between allele frequencies in the two replicates across the genome. We initially observed the segregation distortion in progeny from the NF54-GFPLuc x NHP4026 cross replicate. The major peak on chromosome 13 coincided with the insertion of the GFP cassette in to the *pf47* locus in the NF54-GFPLuc parasite which we hypothesized could be the reason for the distortion. Therefore, we repeated the NF54WT x NHP4026 cross using the unedited parental NF54 with NHP4026 to test if the genetic modification was the driver of the distortion. This was not the case and the repeatability of the skews strongly supports the alternative hypothesis that the GFP cassette is not the driver of this distortion, allowing us to combine the progeny from NF54 x NHP4026 in estimating genetic maps.

### Distorted Loci

We examined each distorted locus for plausible driver genes.

Chr7: a region of 520 kb on chromosome 7 containing 121 genes showed significant segregation distortion in both biological replicates of NF54 x NHP4026 (chi squared test, p<0.001, Fig 4B, S5 Table). This region is disproportionally inherited from NF54 with the most highly distorted region having only 0.05% NHP4026 alleles in NF54GFPLuc x NHP4026 replicate and 0% NHP4026 in the NF54 x NHP4026 replicate. This highly distorted region contains 17 genes (Fig 4B) including *pfcrt* (*PF3D7_0709000*). NHP4026, along with three recombinant progeny, are each highly resistant to chloroquine *in vitro*. Mutations in *pfcrt* are the main driver of chloroquine resistance and have been shown to confer a fitness costs in some genetic backgrounds [31].

**Fig 4.**
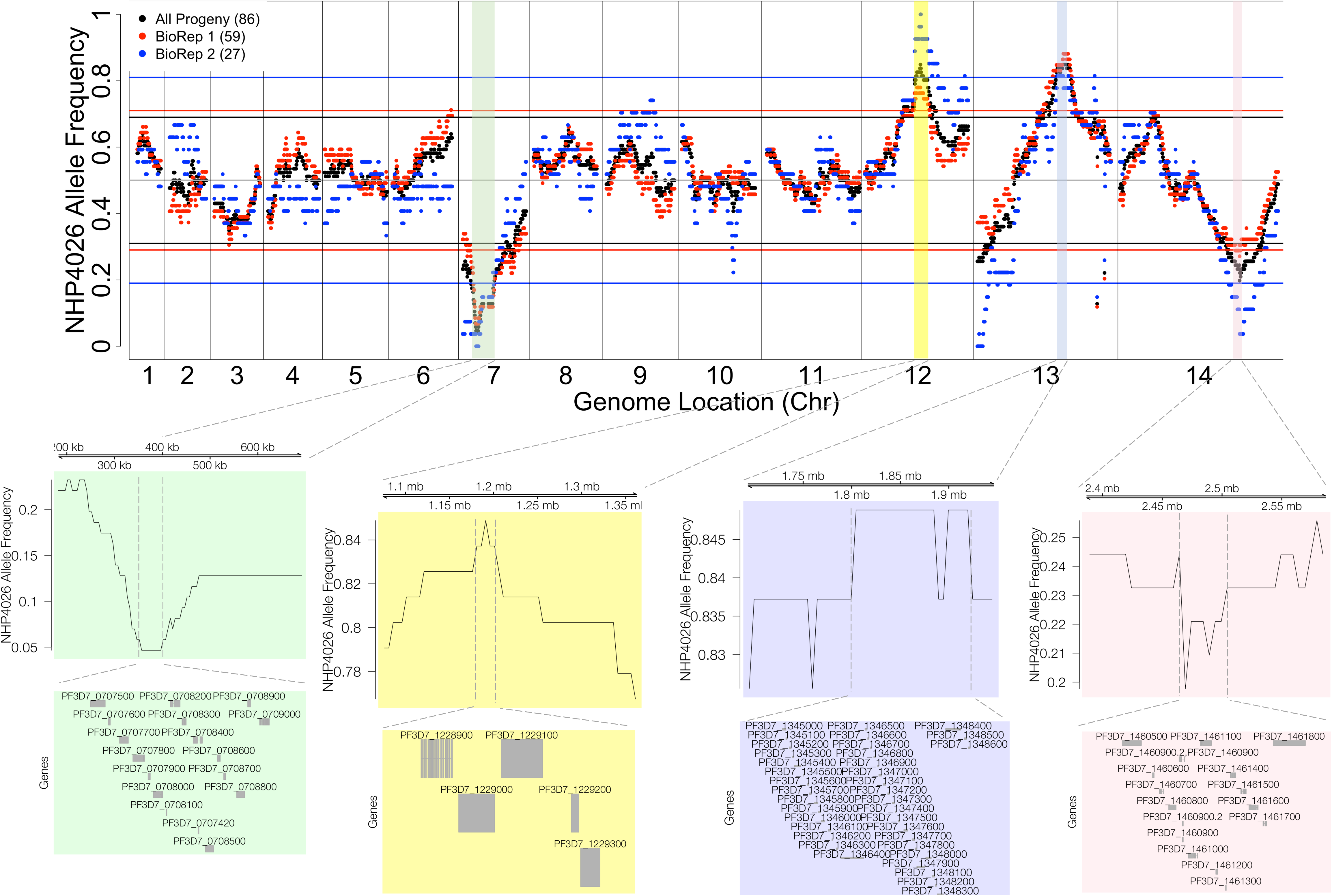
Segregation Distortion Decreases Power and Mapping Resolution. (A) Frequency of the NHP4026 SNP alleles in unique recombinant progeny in NF54 x NHP4026 is highly repeatable across biological replicates (black – all progeny, red – progeny from biological replicate 1, blue – progeny from biological replicate 2). Horizontal lines represent significance thresholds (chi sq p=0.001) for segregation distortion for each corresponding set of progeny. Colored regions show significant segregations distortion in both biological replicates. Genes are shown for the most highly skewed sub-regions.

Chr12: a 295 kb region (with 71 genes) shows replicated significant segregation distortion with an overabundance of NHP4026 alleles. The most skewed region contains five genes (Fig 4C) including *pfmrp2* (*PF3D7_1229100*) at the center of the peak. *Pfmrp2* has high genetic variability among Thai clinical isolates with single genetic variants having significant associations with *in vitro* response to chloroquine, mefloquine and piperaquine and *in vivo* parasite clearance [32].

Chr13: a 230 kb region predominantly inherited from NHP4026 with 56 genes that shows replicated significant segregation distortion. The most highly distorted subregion on chromosome 13 contains *pf47* (*PF3D7_1346800*). In the NF54GFPLuc x NHP4026 replicate of this cross the NF54 line contained a GFPLuc cassette inserted in *pf47* [23] however this insert is not present in the NF54 parent used in the NF54WT x NHP4026 replicate of this cross, which shows the same distortion pattern.

Chr14: a 205 kb region containing 62 genes on chromosome 14 showed replicated significant segregation distortion with alleles predominantly inherited from NF54. The most highly skewed sub-region contains 15 genes including *pfarps10* (*PF3D7_1460900*) and has been associated with slow clearance in GWAS studies [7] and is hypothesized to contribute to a permissive background for evolution of *pfk13* mutations.

Previous *P. falciparum* genetic crosses exhibited significant segregation distortion at several loci [13, 16, 17, 25]. We explored overlap between distorted regions in all the published *P. falciparum* crosses and our two new crosses and included a previously published bulk analysis of selection in uncloned progeny of the MKK2835xNHP1337 cross [33] (S2 Fig). We observe overlaps on chromsomes 12, 13 and 14.

### Increased mapping power in an expanded genetic cross

Previous genetic crosses have been used to map the genetic basis of a wide range of traits. However, small sample size (Table 1) and rampant segregation distortion (S2 Fig) have likely limited detection to mutations with very large effect size (ES). A quantitative dissection of this has not been performed for malaria parasite crosses. To quantify the extent to which our expanded progeny set will improve genetic mapping for the malaria community we performed extensive simulations. We quantified the impact of phenotypic replication, progeny number and the number of loci determining a trait to the power to map a trait and mapping resolution using the 84 progeny from NF54 x NHP4026 (Fig 5 and S3 Fig). Briefly, we used the full progeny panel from NF54 x NHP4026 (n = 84) or subsamples of this panel (n = 60, 50, 40, 30) and simulated phenotypes at different effect sizes using loci with balanced inheritance (0.5 allele frequency) to simulate the phenotype (see Methods for details). We then determined whether the phenotype mapped to the correct loci with a significant LOD score (true positive), did not have a significant association (false negative) or mapped to a different locus (false positive). Using progeny panels comparable to previous genetic crosses (n = 30-40) only very large effect sizes (ES > 0.5) can be mapped with high power (>80%). In contrast, 84 progeny enable mapping of much smaller effect sizes (ES = 0.2) at 80% power. Increasing the number of progeny also increases the locus resolution (S3 Fig). At an ES of 0.5, with n = 30 we can on average map to a region containing 58 genes; moreover, with n = 84, we can map to a region containing only 17 genes (S3 Fig). At small effect sizes we observe similar large increases in mapping resolution as we increase the size of the progeny set and more modest increases for larger effect sizes (S3 Fig).

**Fig 5.**
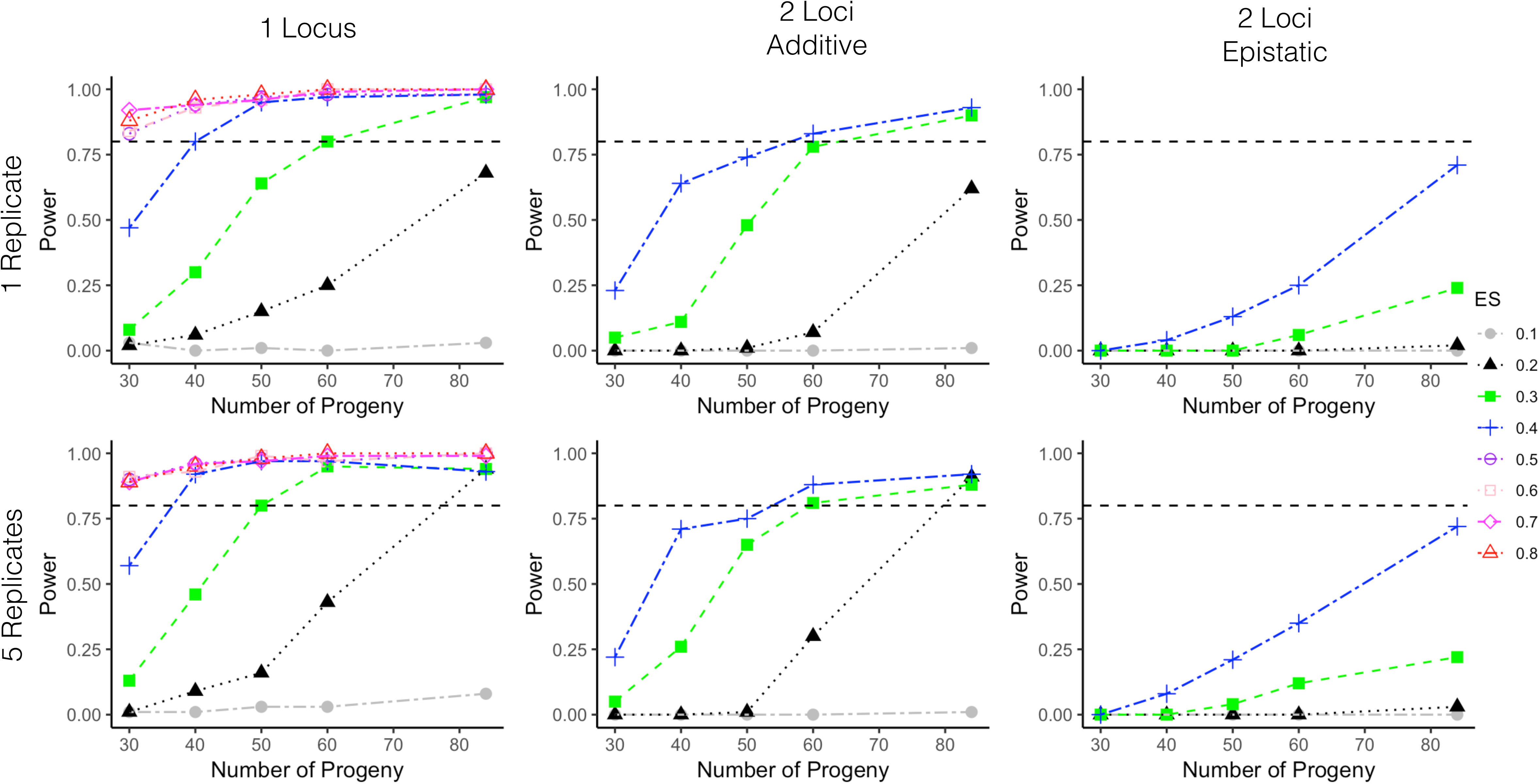
Power analysis for different size progeny sets. Power curves are shown from simulated phenotypes for NF54 x NHP4026 progeny for different size progeny sets. The top row shows power curves where the phenotype only has a single replicate per progeny strain and the bottom row shows results for 5 replicate phenotype values per progeny strain. The first column shows results for a single locus effect, the second column shows results for an additive 2 loci effect and the third column shows results for an epistatic interaction between 2 loci. The horizontal black line denotes 80% power.

Most genetic traits are not monogenic but are complex in nature. To better capture the complex genetic architecture, multiple loci must be identified and these loci sometimes interact (i.e. do not contribute individually and additively). For a trait controlled by two additive loci that contribute equally to the phenotype, 60 progeny, with replicated phenotypes can detect an association at ES = 0.3, whereas 84 progeny are needed to detect an association at ES = 0.2. When a trait is controlled by two epistatically interacting loci, 84 progeny with replicated phenotypes provide 75% power to detect an association and interaction with ES = 0.4. Replicated phenotypes allow the same power to be achieved with fewer progeny for ES ≥ 0.3 and allow for a trait with ES = 0.2 to be detected for N = 84 progeny for additive loci. This analysis indicates that the four previous *P. falciparum* crosses (conducted in chimpanzee hosts) generating from 15 - 35 progeny, were underpowered. In progeny sets of this size could reliably detect associations only for phenotypes with large effects sizes, ES ≥ 0.5 and were not able to detect even a very strong epistatic interaction. Our two new crosses with n = 60 and 84 progeny have much higher power and are capable of reliably detecting phenotypes with effect sizes as low as 0.2.

Polygenic traits don’t always have equal contributions from multiple loci. In malaria parasites, there are several well-known phenotypes with one known major effect locus [34], including chloroquine resistance and mutations in *pfcrt*, sulfadoxine and point mutations in *pfdhps*, pyrimethamine and point mutations in *pfdhfr*, atovaquone and point mutation in *pfcytb*, mefloquine and *pfmdr1* and artemisinin resistance and mutations in *pfk13.* It is an open question whether we could detect more subtle secondary loci with genetic crosses with additional progeny. Our analysis (Fig 6) shows that with 84 progeny we can detect secondary loci with ES as low as 0.2 and 0.15. However, with smaller numbers of progeny we are not able to detect both contributing loci. With 60 progeny we can detect only one of the secondary loci at ES=0.2 and with 35 progeny we can detect neither secondary loci.

**Fig 6.**
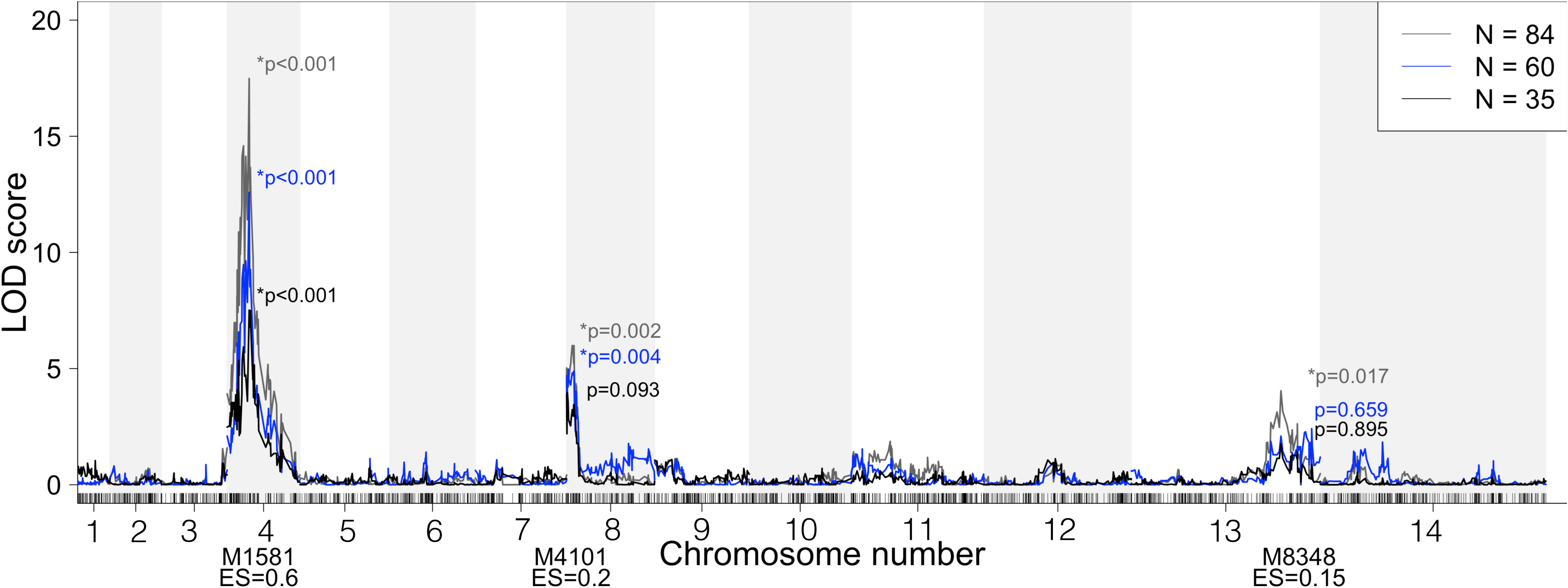
Detecting complex associations. QTL scans of simulated phenotypes with one major (ES=0.6) and two minor (ES=0.2 and 0.15) contributing loci for N=84 (grey), 60 (blue) and 35 progeny (black). The major locus is detected for all sizes of N, but only one minor locus is detected for N=60 progeny and neither minor locus is detected at N=35 progeny.

### Segregation distortion decreases the resolution and power of mapping

Segregation distortion is abundant across nearly all *P. falciparum* genetic crosses generated to date, with our newly generated MKK2835 x NHP1337 cross being the sole exception. We performed a power analysis to determine the impact of segregation distortion on the power to identify causal variants. Segregation distortion decreases power to detect effects near the distorted locus, especially for phenotypes with small effect sizes (Fig 7). For phenotypes with large effect sizes and for large numbers of progeny, the extent of segregation distortion in the F_1_ mapping population at the controlling locus has little effect on power; however, as the number of progeny decrease, a significant loss of power occurs as the degree of segregation distortion increases. The loss of power due to segregation distortion is even more pronounced with fewer progeny (Fig 7). For an ES of 0.8, we can detect associations for loci with any allele frequency using as little as 30 progeny. For an effect size of 0.4, 50 progeny are necessary to detect an association for allele frequencies ranging from 0.3 to 0.7, and only 84 progeny will allow us to detect an association at a more distorted loci with 0.2 or 0.8 allele frequency. At 0.2 effect size we can only reliably detect an association for a locus with even segregation using 84 progeny.

**Fig 7.**
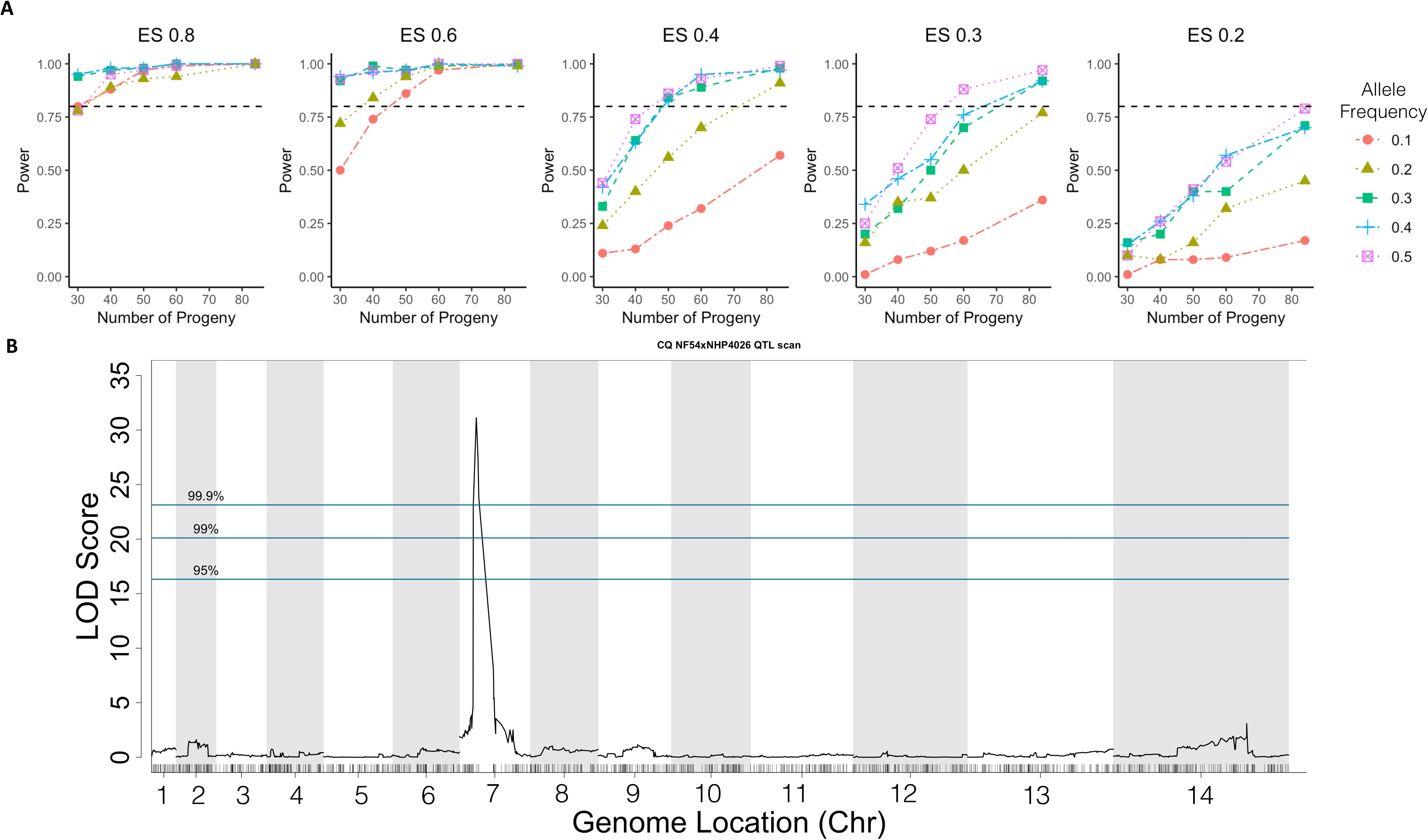
Power loss due to segregation distortion. (A) Effect of SD on mapping power in NF54 x NHP4026 with simulated phenotype data at different effect sizes. Each sub-panel shows the relationship between allele frequency and power for different numbers of progeny at a fixed effect size. For high effect size, allele frequency has little effect on power. At lower effect sizes we observe a large loss of power for alleles with less than 0.3 allele frequency. (B) QTL mapping of CQ IC_50_ (ES=0.84) in 35 progeny in the NF54xNHP4026 cross results in a LOD score of 18 and a genome wide p-value = 0.000696 showing that in real data with extreme SD a trait with high effect size is detectable.

In NF54 x NHP4026 we observe significant segregation distortion (p<0.001) with allele frequencies at distorted loci ranging from 0.05 to 0.31 and 0.69 to 0.85 (Fig 3), including in regions that include important drug resistance genes including *pfcrt* (Chromosome 7) and *pfk13.* Despite this extreme segregation distortion on chromosome 7 in NF54 x NHP4026 (NHP4026 allele frequency of 0.05), it is still possible to map the chloroquine drug response to the locus containing *pfcrt* (p<0.00001, Fig 7B). In contrast, in MKK2835 x NHP1337, allele frequencies of the NHP1337 alleles range from 0.3 to 0.7 (Fig 3). At these allele frequencies we see consistent power indicating that power and mapping resolution are expected to be consistent across the genome.

## Discussion

### Power of *P. falciparum* genetic crosses generated using FRG NOD huHep/huRBC mice

Historical challenges to generating novel *P. falciparum* genetic crosses made GWAS, *in vitro* selection experiments and bulk sequencing approaches the more effective means to study new drug resistance-associated phenotypes as they emerge in the clinic. However, each of these techniques has its limitations. *In vitro* selections are time consuming, sometimes requiring several years to produce resistant lines, and may not identify loci evolving under drug pressure in the field situation [35]. GWAS is often confounded by population structure and has low power to dissect complex genetic traits, i.e. multiple loci, multiple alleles per locus and epistasis [36]. On the other hand, a well-conceived and controlled genetic cross can greatly complement these techniques, as each cross can be designed to answer specific questions and then have high power to dissect complex associations between genotype and phenotype. Historically, *P. falciparum* controlled genetic crosses have been made with splenectomized chimpanzees strictly limiting their production due to cost and ethical concerns. Use of the human tissue-chimeric FRG NOD huHep/huRBC mouse restores and expands our ability to make controlled genetic crosses in malaria parasites [23]. We demonstrate here that targeted crosses between clinical isolates can be generated in real time (six months) and outperform all previous crosses in their size, mapping power and precision.

We created the first *P. falciparum* cross between two sympatric recent clinical isolates from the Thai-Myanmar border, MKK2835 and NHP1337. Analysis of the recombination rate, segregation distortion, and selfing rate of this cross revealed interesting differences to all other *P. falciparum* crosses including our NF54 x NHP4026 cross, between a lab line and a recent field isolate. In MKK2835 x NHP1337 we observed minimal segregation distortion and a high percentage of clones that resulted from selfing. We have also shown that most of the recombinant progeny recovered are unique when cloning is initiated immediately, or within five days of establishing *in vitro* culture. Using simulations, we have shown that the power to detect associations between phenotypes and genotypes increases drastically when we are able to map with populations with 60 – 84 individuals. We have also shown through simulation and using real phenotype data that segregation distortion can lower power to detect QTL at distorted loci even for phenotypes with moderate effect sizes. Nevertheless, major effect loci can be mapped within these regions, supporting the utility of our crosses in these cases. Furthermore, because we can cryopreserve uncloned F_1_ parasite populations, it is possible to further isolate additional independent recombinant progeny for future analyses, as sequential cloning attempts will isolate new unique progeny.

### Power of malaria crosses generated using humanized mice

We have shown that the FRG NOD huHep/huRBC mouse can be used to rapidly make controlled genetic crosses on demand from field isolates to create F1 progeny populations with large numbers of clonal recombinant progeny per cross. This dramatically increases our power to detect associations with greater resolution. Using new crosses with more recombinant progeny and higher power we can dissect genetic architecture and determine the individual contributions of different loci to polygenic traits. We can also map phenotypes with small to modest effect sizes more precisely, to smaller regions of the genome. For instance, at an ES of 0.5 using 30 recombinant progeny, we can map to a region containing 58 candidate genes. However, at an ES of 0.5 using 84 progeny, we can map to a region of 17 candidate genes (S3 Fig). With increased transfection efficiencies using CRISPR/Cas9-based technology, it is not unreasonable to then target the genome by transgenesis to pinpoint loci involved in observed phenotypes. For phenotypes with large ES, similar to that conferred by chloroquine resistance (0.8), with 30 progeny we can map to a region containing on average 20 candidate genes whereas with 84 progeny we can map to a region containing only eight genes. These significant reductions in number of candidate genes has a large impact on our ability to determine causal mutations, drastically reducing the effort required for validation studies. Furthermore, our ability to generate further genetic crosses between the same two parents of interest is unparalleled, allowing us to potentially isolate 100’s of unique recombinant progeny for analysis.

### Maximizing Numbers of Unique Recombinant Progeny

Based on the prevalence of infected mosquitos and estimates of oocysts/mosquito we can estimate the number of unique recombinants in the mosquitos used to infect each FRG NOD huHep/huRBC mouse. During the parasite lifecycle there are multiple bottlenecks which reduce the number of genotypes in a blood stream infection. Oocysts may arise due to selfing or fail to progress, sporozoites may fail to reach the liver and further attrition through the liver and blood stages will occur. As we observed, without extensive cloning efforts we are unable to capture all these possible unique recombinant progeny. Interestingly, each cloning round produced almost entirely unique sets of progeny indicating that our cloning efforts (166 clones for NF54 x NHP4026 and 266 for MKK2835 x NHP1337) under-sampled the total population of recombinant progeny available. Recovering unique sets of progeny from each cloning round indicates that there are likely many more additional unique progeny to recover from the bulk F_1_ populations and that strategic additional cloning would likely provide a substantial return in F_1_ progeny numbers. In order to maximize the number of unique recombinant progeny recovered, we showed that cloning straight after the *in vivo* liver stage to blood stage transition or as early as possible after establishing *in vitro* culture gave a large degree of success. Also, initiating cloning either directly from the mouse or from a thawed stock of bulk culture did not impact the proportion of unique recombinant progeny recovered. Notably, we minimized the potential for additional loss in diversity during cryopreservation by freezing immediately after exsanguination and cloning within 48 hours of thaw. Additionally, with streamlining of the crossing process and being able to complete a cross from thawing of parental lines to isolating, genotyping and identifying unique recombinant progeny within 6 months it is easy to simply repeat the cross and generate an entirely distinct set of recombinant progeny to generate additional unique recombinant progeny.

### Differences in selfing between crosses

*P. falciparum* infections in nature are sometimes monoclonal and sometimes co-infections, depending on the genetic diversity of the gametocytes taken up during a mosquito blood meal. Thus, *P. falciparum* must be able to maintain its life-cycle through selfing as well as out-crossing. Evidence from natural infections suggests that mating can by non-random when distinct parasite lineages are co-transmitted from a single mosquito bite [37]. In previous crosses between established lab lines 3D7 and HB3, it was shown that selfed progeny are observed at expected ratios in oocysts [38, 39] and early in blood stage culture, but at lower than expected ratios among clones when cloning was begun 32 days after isolation from chimpanzees [40, 41]. In 7G8 x GB4, 29 of more than 200 (14.5%) individual clones were selfed [25].

Our MKK2835 x NHP1337 cross between two recent field isolates, both from Southeast Asia, produced more selfed progeny than previously reported for *P. falciparum* genetic crosses. Interestingly, NHP1337 dominated the selfed progeny almost entirely, consistent with bulk allele frequencies in samples taken at similar times [33]. While efforts were made to infect the mosquitos with equal number of MKK2835 x NHP1337 gametocytes, the unequal selfing rates may reflect an imbalance in the initial gametocyte ratio or in gametocyte viability between MKK2835 and NHP1337. It is also possible that there are inherent difference in selfing rates between MKK2835 and NHP1337, although both lines successfully selfed in mosquito cages infected with only one parent (S1 Table). We do not yet know if the large proportion of selfed clones observed in our cross between recent field isolates will be repeated in future crosses. Using bulk segregant analysis of these same populations we suspect that these selfed clones are outcompeted over time in non-stressed *in vitro* culture conditions [33] this may perhaps also have been the case in the previous 3D7 x HB3 cross [39, 40].

In contrast we observed very few selfed progeny in our NF54 x NHP4026 cross, between a recent Southeast Asian field isolate NHP4026 and the established African lab line NF54. Both NF54 and NHP4026 readily self when in used alone to inoculate mosquito cages with NF54 often giving very high infection prevalence and numbers of oocysts/midgut (S1 Table). In several cloning rounds of NF54 x NHP4026, cloning was initiated immediately after transition to *in vitro* culture, indicating that in this cross selfed progeny were not selected against in bulk competition with recombinant progeny. Further experiments will be necessary to understand why NF54 x NHP4026 generated so few selfed progeny.

### Differences in segregation distortion between crosses

In other systems segregation distortion is often more extreme when more distantly related parents are crossed. For instance, interspecific crosses have been shown to result in segregation distortion more often and with more severe distortion that intraspecific crosses [42, 43]. All previous *P. falciparum* genetic crosses were between allopatric parasite lines, generally isolated on different continents, and unsurprisingly show significant segregation distortion over large regions of the genome. Similar to previous *P. falciparum* crosses, our allopatric cross between an establish lab line (NF54 or NF54GFPLuc, African origin) and a recent field isolate (NHP4026, Thai-Myanmar border) had regions of significant segregation distortion that were consistent across replicates. Conversely, the MKK2835 x NHP1337 cross which relied on two sympatric parasites recently isolated from the Thai-Myanmar border was the first *P. falciparum* controlled genetic cross to have relatively even inheritance patterns across the genome with no significant segregation distortion. One possible explanation for the observed segregation distortion is that natural selection may act against unfit allele combinations causing a deviation from expected mendelian rations [44]. It is also possible that there are prezygotic barriers such as barriers to gamete recognition between more distantly related parents.

In NF54 x NHP4026, the subregions with the most highly skewed allele frequencies in each of the significantly distorted regions contain genes of interest. The most highly distorted subregion on chromosome 7 (predominantly inherited from the NF54 parent with only 3 progeny inheriting alleles from NHP4026) includes *pfcrt* which is known to carry a substantial fitness cost in some genetic backgrounds and that different combinations of mutations are more deleterious than others [31]. Although NHP4026 is a parasite that grows particularly well in *in vitro* culture [45] (even outcompeting NF54 in co-culture experiments) it is clear that inheriting an NHP4026 allele at this locus contributes a fitness cost. The most highly skewed subregion on chromosome 14 is also predominantly inherited from NF54 and contains *pfarps2* which has been associated with artemisinin resistance (slow clearance of parasite from treated patients) in GWAS studies and is thought to contribute to a permission background for development of artemisinin resistance [7]. While NHP4026 is *pfk13* WT it does have a slow clearance phenotype. It will be interesting to explore weather *pfarps10* has a fitness cost in this genetic background. Interestingly, while we see no segregation distortion in MKK2835 x NHP1337 among the cloned progeny, we do see selection on chromosome 14 over time in a uncloned bulk culture of MKK2835 x NHP1337 cross F_1_ progeny that is also centered on *pfarps10* where selection is against the derived alleles in *pfarps10* [33].

Alternatively, on chromosomes 12 and 13 there are subregions where alleles are more commonly inherited from NHP4026. The region on chromosome 12 include *pfmrp2* and the region on chromosome 13 includes *pf47*. The most skewed region on chromosome 12 overlaps with the selected region in a uncloned bulk culture of MKK2835 x NHP1337 cross F_1_ progeny except that selection is against the derived allele in *pfmrp2* in this case. *Pfmrp2* has been associated with mefloquine and piperaquine response *in vitro* and parasite clearance [32] in Thai isolates and we speculate it may have a fitness cost *in vitro.* The role *pfmrp2* plays in drug resistance is still unclear and these genetic crosses may help elucidate its function. In the NF54GFPLux x NHP4026 replicate, the NF54 parent contained a gfp/luciferase cassette insert on chromosome 13 in *pf47*, the NF54WT x NHP4026 replicate of this cross was made with the isogenic NF54 line without the gfp/luciferase insert. We saw consistent inheritance patterns in both biological replicates of this cross indicating the skew here is gfp/luciferase insert independent. A large region of segregation distortion was observed on chromosome 13 in 7G8 x GB4, part of which overlaps our region of segregation distortion in NF54 x NHP4026 [17, 25]. *Pf47* and *pfs45/48*, two 6-cys proteins are located in the center of this subregion. These two genes are known to by highly polymorphic in natural populations and are thought to be under selection because of roles in gamete recognition and compatibility [46, 47]. It is possible that *pf47* and/or *pfs45/48* play a key role in segregation distortion in more distantly related lines but not in a cross between allopatric recent clinical isolates. Indeed, we observed no significant segregation distortion in the MKK2835 cross and also observed no selection over time on chromosome 13 in the bulk segregant experiment using the MKK2835 x NHP1337 bulk F_1_ progeny [33].

We think that natural selection acting against unfit allele combinations is a plausible explanation for some regions of segregation distortion in NF54 x NHP4026 including the regions on chromosome 7, 12 and 14 and the observed selection over time in the uncloned bulk F_1_ culture from the MKK2835 x NHP1337 cross [33]. Issues with gamete recognition and compatibility might drive segregation distortion observed on chromosome 13 in NF54 x NHP4026 and 7G8 x GB4 (both allopatric) but not in the sympatric MKK2835 x NHP1337 cross (see also [33]). Performing competition experiments between individual progeny with different alleles at these distorted and selected loci will be informative in determining how different combinations of alleles might contribute to parasite fitness [45]. These experiments can be followed with CRISPR/Cas9 editing of polymorphisms in individual genes as further validation.

### Loss of power at segregation distortion loci

Segregation distortion loci traditionally have been excluded in genetic mapping studies to avoid loss of power to detect real effects (type II error, false negative) and the potential to detect false positives (type I error) [48]. Excluding distorted loci from analysis would be particularly problematic in *P. falciparum* because all previous crosses had large regions of significant segregation distortion that contain known resistance loci. Using segregation distortion loci in mapping studies is possible, however it is necessary to carefully interpret results keeping in mind the loss of power to detect effects in distorted regions. If drug resistance loci are at or near genome regions showing segregation distortion loci, we may fail to detect these drug resistance loci in crosses with small numbers of progeny or when effect size is small. We have shown this effect through mapping simulated phenotypes to loci with varying degrees of segregation distortion. Despite the extreme segregation distortion observed in NF54 x NHP4026 (NHP4026 allele frequency of less than 0.05 at *pfcrt*) and only three progeny plus NHP4026 showing a chloroquine resistant phenotype we are able to correctly map the chloroquine drug response to the locus containing *pfcrt*. Through simulation, we demonstrate reliable detection of QTL for phenotypes with very large effect sizes (ES = 0.8), even for very distorted loci and small numbers of progeny. However, as the effect size decreases, we observe stronger loss of power at distorted loci. For phenotypes with moderate effect sizes we can only reliably detect QTL at distorted loci using large numbers of progeny. Therefore, care is required in interpreting negative QTL results for phenotypes with small to moderate effect sizes, especially when mapping in small progeny sets. When QTL and segregation distortion loci coincide, false negatives will lead us to miss real associations between phenotypes and genetic variants. This problem with power will be amplified when attempting to map omics phenotypes where multiple testing correction must be employed. However, while segregation distortion presents a challenge for linkage analysis, the location of genome regions showing strong skews can help to pinpoint loci with large phenotypic effects.

## Conclusions

We believe that the use of the human hepatoycte-chimeric FRG NOD huHep/huRBC mouse to generate genetic crosses in *P. falciparum* has the potential to revolutionize quantitative genetics in *P. falciparum*. It is feasible to generate crosses on demand to study the genetic architecture of emerging phenotypes. We can also use complex cross designs to improve power to detect associations for phenotypes where a genetic variant only controls a small amount of variation. Shared parent crosses are ideal for understanding the role of individual mutations within phenotypes with complex genetic architecture. Pairwise crosses of a small group of isolates can be used to create a diversity panel that captures a large amount of phenotypic and genetic variation in *P. falciparum*. Similarly, many other complex cross designs that have been used extensively in the plant and animal breeding literature that are now open to malaria researchers.

## Methods

Genetic crosses were conducted largely as described previously [23]. We made several adjustments to maximize recovery of progeny from the genetic crosses, including completing independent replicates of the crosses and cloning via limiting dilution directly from the transitioned blood removed from the FRG NOD huHep mouse. In addition, the transition to *in vitro* culture was carried out using media containing Albumax rather than human serum. We observed successful expansion of the transitioned cultures in both serum-containing and Albumax-containing media, but downstream limiting dilution cloning failed to yield the expected number of clones if carried out using serum. We therefore cloned and expanded the transitioned blood stage culture in media containing Albumax. Screening for clones was carried out using the Phusion Blood Direct PCR Kit (Thermo Scientific). Specific methodological information for each replicate of each cross is provided in S2 Table.

### Identifying Positive Clones

Beginning at week 2 post cloning and continuing until week 6 the Phusion Blood Direct PCR Kit (Thermo Scientific) was utilized to identify positive clones. This kit is very sensitive, detecting positive parasitemia using only 1 μL of infected culture streamlining our detection of positive clones. A protocol for this screening method is available in the S1 File.

### MS Genotyping

All progeny of NF54 x NHP4026 were initially genotyped via microsatellite markers to identify unique recombinants. The progeny isolated in cloning rounds 1 and 2 or replicate 1 of the NF54 x NHP4026 were genotyped at 17 MS markers. The progeny isolated in cloning round 3 of NF54 x NHP4026 were genotyped at 8 MS markers. Primers for each MS marker used are listed in S6 Table. For cloning rounds 1 and 2, full genome sequencing was performed for all unique recombinants. For cloning round 3 and all other crosses all potential recombinant progeny were fully sequenced.

### Preparation and sequencing of progeny

DNA was extracted from 35-50uL of packed red blood cells using Quick DNA Kit (Zymo). Libraries were prepared with ¼ reaction volumes of the KAPA HyperPlus DNA Library Kit and 20-50ng of extracted DNA according to manufacturer directions with slight modifications. Fragmentation time was 26 minutes; adapter ligation was increased to 1 hour; PCR was performed for 7 cycles; and size selection was performed post PCR using full volume methods. We used KAPA Dual-Indexed Adapter Kit, adding 7.5uM adapter to the appropriate well. Samples were measured for DNA quantity using the QBit BR DNA Kit. Samples were then pooled for sequencing based on their QBit measurements to normalize input. The pooled sample was quantified using the KAPA Library Quantification Kit, and adjusted to 2-4nM with 10mM Tris-HCl, pH 7.5-8.0 (Qiagen) for sequencing on Illumina platforms. The pool was also run on the Agilent Tape Station using the D1000 BR Kit to assess sample size and lack of primer dimers. Pools were run on the Illumina HiSeq 2500 or Illumina NextSeq for 2×100bp run

We aligned raw sequencing reads to v3 of the 3D7 genome reference (http://www.plasmodb.org) using BWA MEM v0.7.5a [49]. After removing PCR duplicates and reads mapping to the ends of chromosomes (Picard v1.56) we recalibrated base quality scores, realigned around indels and called genotypes using GATK v3.5 [50] in the GenotypeGVCFs mode using QualByDepth, FisherStrand, StrandOddsRatio VariantType, GC Content and max_alterate_alleles set to 6. We recalibrated quality scores and calculated VQSLOD scores using SNP calls conforming to Mendelian inheritance in previous genetic crosses, and excluding sites in highly error-prone genomic regions (calls outside of the “core genome” [21].

### Filtering high quality SNP variants

The .vcf file containing parents, potential progeny and all high quality SNPs were processed in R using the vcfR library. Initially SNP filters were based on the parental distributions; only homozygous, bi-allelic parental SNPs with high coverage (≥ 10) and high quality scores (GQ ≥ 99) were retained. Next, low quality SNPs across parents and progeny were filtered with a VQSLOD < 2.5. This final SNP set was defined as our high quality SNP set for further analysis.

### Filtering Progeny

In *P. falciparum* crosses to produce the F_1_ mapping population, it is necessary to filter out potential progeny that are non-clonal and repeated sampling of the same genotype. Initially, potential progeny with more than 80% missing data were removed from further analysis.

### Identifying and filtering non-clonal progeny

Since *P. falciparum* parasites are haploid throughout the entirety of the human portion of their life-cycle including the intraerythrocytic stage during which they are cloned we expect that clonal infections should have predominantly homozygous SNP calls except for rare instances of sequencing error. In contrast, non-clonal infections where the mixture contains full siblings or full siblings and parent genotypes would have contiguous regions with high numbers of heterozygous SNP calls at above the rate expected from sequencing error along.

The sequencing error rate was estimated for each cross as the mean from a distribution of percent heterozygous SNP calls across all potential progeny (S4 Fig). Assuming true sequencing errors follow a Poisson process with λ = % sequencing error, then the expected distance between sequencing events as ^1^/_λ_. To identify non-clonal samples we counted heterozygous SNP calls across the genome in a sliding window of size ^1^/_λ_ and using a Poisson Distribution with λ = % sequencing error calculated the probability of getting at least the observed number of heterozygous SNP calls in each window. These probabilities were adjusted for multiple testing based on the number of windows in the genome and the adjusted probabilities were plotted as a heatmap (S4 Fig). Samples with windows with adjusted probabilities < 0.05 were designated as non-clonal and filtered from the final progeny set.

### Phasing of clonal progeny

A matrix of phased genotypes was constructed for parents and clonal progeny for all high quality SNPs. In each cross the drug sensitive parents (NF54 and MKK2835) were coded as 0 while the drug resistant parent (NHP4026 and NHP1337) were coded as 1. Progeny SNPs that matched the drug sensitive parent’s SNPs were coded as 0 while SNPs that matched the drug resistant parent’s SNPs were coded as 1. Heterozygous SNPs were coded as missing.

### Identifying Unique Recombinants

Our high quality phased dataset for clonal progeny was formatted for the qtl package in R and loaded as a genetic map. Genotype similarity scores were computed using the comparegeno function. A similarity score cut-off of 0.9 was used to define clusters of genetically distinct recombinant progeny (see S1 File for details of cut-off was chosen). Individual progeny were selected from each cluster of genetically similar progeny using igraph in R. Only unique recombinant progeny and parents were retained to create a final dataset of all SNPs.

### Physical Recombination Map Construction

5kb windows were defined across the core genome to construct a heatmap depicting a physical recombination map for each cross. For each progeny, in each 5kb window the most common parental genotype was determined, if a window contained only missing data then it was filled if the next window with data had a matching genotype to the previous window with data, otherwise it was left missing.

### Defining informative markers

All phased genotype data for clonal, unique recombinant progeny and parents were loaded into R qtl as a genetic map. The findDupMarkers function was used to identify clusters of markers with identical genotype data and the central marker from each cluster was retained in a set of informative markers. This set of informative genotype markers for all clonal, unique recombinant progeny was used for all subsequent analysis and figures. The entire filtering pipeline is available on github (https://github.com/kbuttons/P01_ProgenyCharacterization) with documentation.

### Genetic Map Construction

For each cross the set of informative genotype markers for clonal unique recombinant progeny was coded as A for the sensitive parent and B for the resistant parent and – for missing data and loaded into JoinMapv4.1. Population type was set to HAP1 and the Kosambi mapping function was employed in generating each genetic map. All other parameters were initially set to defaults, however, to account for the systemic segregation distortion observed in the NF54xNHP4026 cross it was necessary to expand the population threshold ranges such that the independence LOD ranged from 1.0 to 15.0, the independence P-value from 1.0e-3 to 1.0e-5, the recombination frequency from 0.250 to 0.001 and the linkage LOD from 3.0 to 15.0. This change in parameters allowed us to differentiate between SNP markers with similar distortion patterns that were known to be physically located on different chromosomes.

### Power Analysis

Progeny from NF54 x NHP4026 were used to estimate power under three different scenarios, one genetic locus contributing to phenotypic variation, 2 loci with additive contributions to phenotypic variation and 2 loci with epistatic interaction controlling phenotypic variation. All models were simulated for the full F_1_ progeny set with N=84 and for subsamples with N=30, 40, 50, 60 and 70. Under the one locus model, a phenotype was simulated as either a single replicate value or the average of 5 replicates at effect sizes ranging from 0.1 to 0.8. Under the two additive loci model, a phenotype was simulated as either a single replicate value or the average of 5 replicates for effect sizes for each locus ranging from 0.1 to 0.4. Under the two epistatic loci model, the first locus controlled whether a trait was present in an on/off fashion and the second locus controlled the level of the phenotype (ie. locus 1 is necessary to be drug resistant and locus 2 controls the level of resistance) and the main effects of both loci ranged between 0.1 to 0.4. A set of markers with 1:1 mendelian inheritance patterns were used as the 1 or 2 loci in the models. All qtl mapping was performed with r qtl. For each simulation, significance thresholds were defined based on 1000 permutations. True positives were defined as a LOD peak that meant the α=0.05 significance threshold and whose 1.5 LOD interval contained the actual marker used in the model.

### SD Power Analysis

This analysis was similar to the 1 locus model in the previous section. In these simulations effect sizes were calculated based on balanced inheritance and levels included 0.2, 0.3, 0.4, 0.6 and 0.8. All markers were categorized by their allele frequency and sorted into bins for each level of allele frequency skew (ie. 0.89 to 0.91 and 0.09 to 0.11 were in the 0.1 bin which represented the most skewed alleles in this analysis). QTL mapping, significance levels and definition of true positives were that same as in the power analysis above.

## Acknowledgements

We would like to thank Jasmine Clark for help with progeny cloning. We would like to acknowledge members of the Ferdig lab, Anderson lab, Cheeseman lab, Vaughan lab, Kappe labs and Emrich lab for helpful discussions.

## Supporting Information

**S1 Fig.**
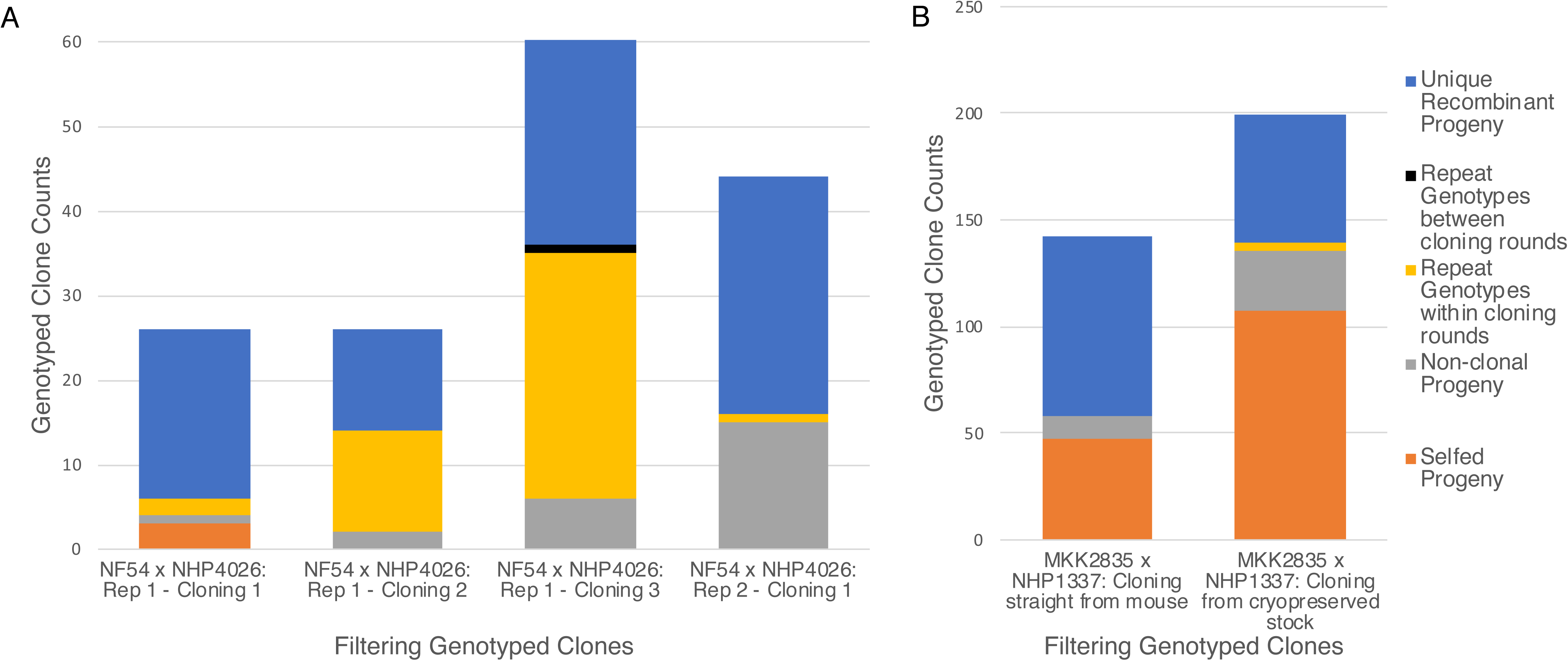
Cloning results for each cross by biological replicate and cloning round. Cloning success varied as a function of length of time parasites were in bulk culture before cloning. (A) Progeny for the NF54 x NHP4026 cross were filtered to identify unique recombinant progeny (blue). Selfed progeny (orange), non-clonal progeny (grey) and repeat sampling of the same genotype within a cloning round (yellow) and between cloning rounds (black) were filtered out of total genotyped progeny for each biological replicate and cloning round. (B) Progeny for the MKK2835 x NHP1337 cross were filtered to identify unique recombinant progeny (blue). Selfed progeny (orange), non-clonal progeny (grey) and repeat sampling of the same genotype within a cloning round (yellow) and between cloning rounds (black) were filtered out of total genotyped progeny for each cloning round.

**S2 Fig.**
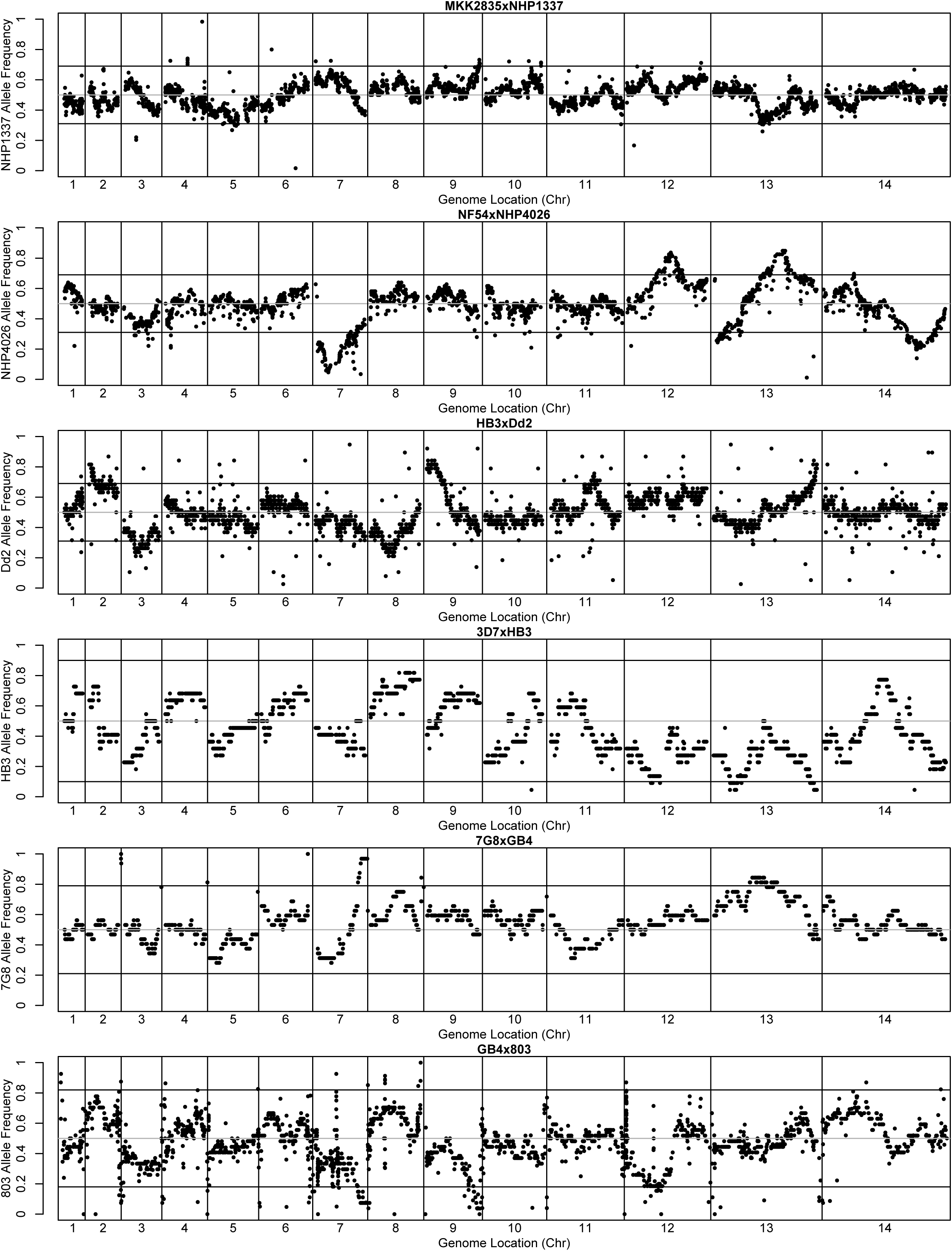
Segregation distortion in all published *P. falciparum* crosses. Allele frequencies plotted across the genome for all 6 published *P. falciparum* crosses show no significant segregation distortion in the MKK2835xNHP1337 cross (A) in contrast to all other published crosses which show regions of significant segregation distortion (B-F).

**S3 Fig.**
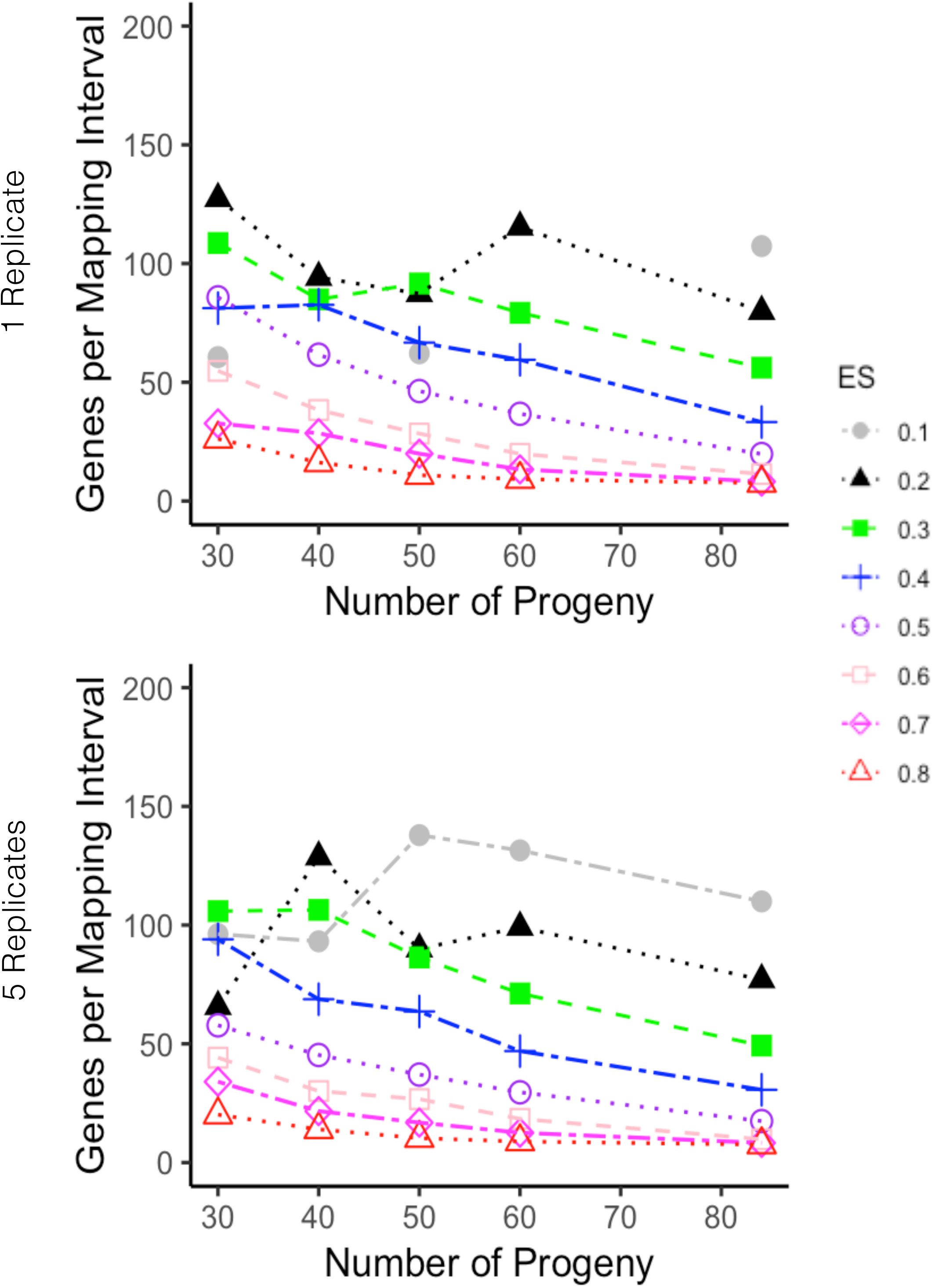
Mapping Resolution for different size progeny sets. Average mapping resolution reported as number of genes per 1.5 LOD interval for simulated phenotypes that accurately map to the 1.5 LOD interval surrounding the causal loci. Progeny set size varied from the full NF54 x NHP4026 progeny set of 84 and was subsampled at 30, 40, 50, 60 and 70 progeny. Each curve represents phenotypes simulated with a given effect size (ES) with ES ranging between 0.1 to 0.8.

**S4 Fig.**
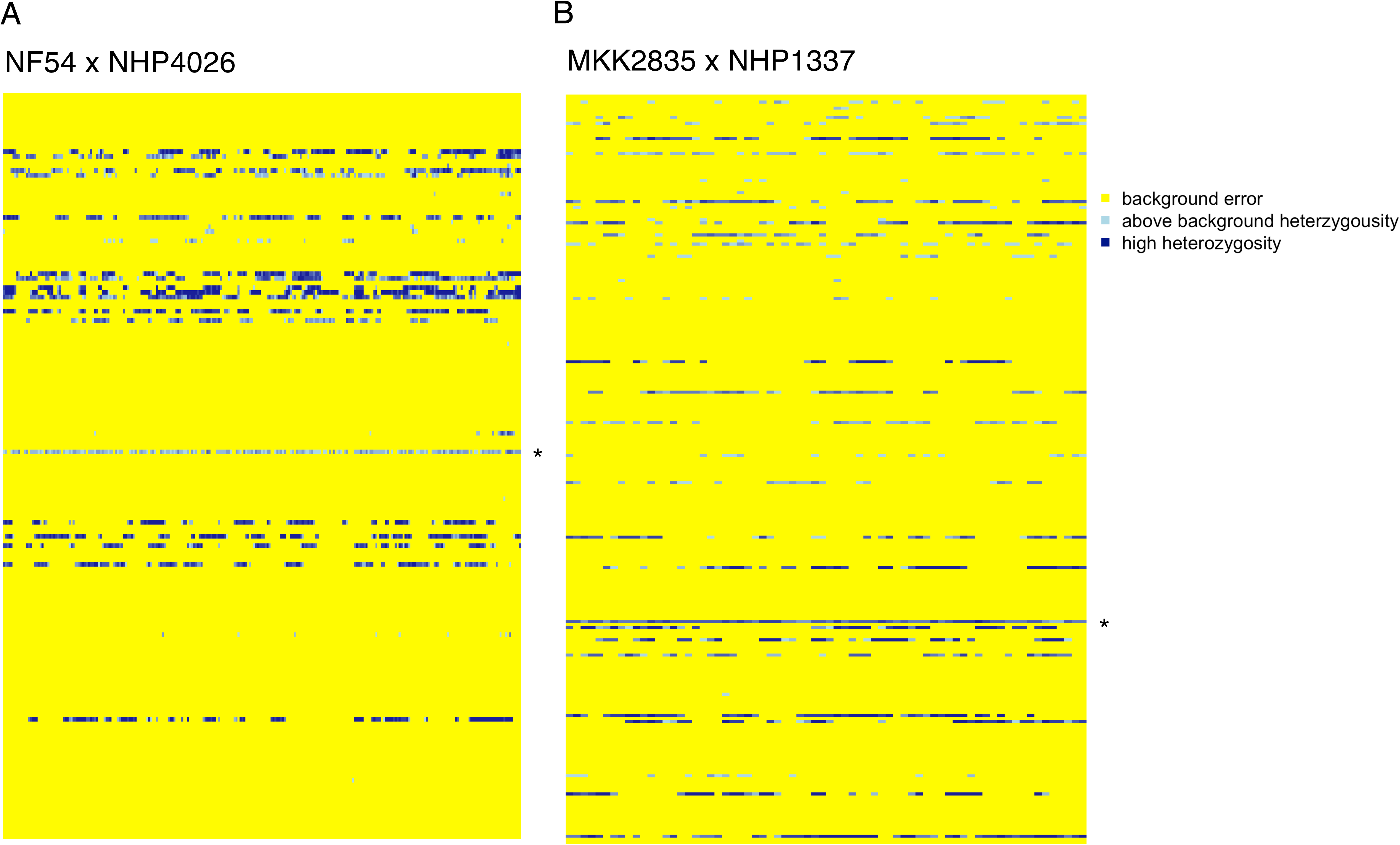
Nonclonal Progeny Heatmap. Heatmap showing regions of the genome for each progeny with above expected numbers of heterozygous allele calls. Regions with above expected heterozygous SNP calls were identified through a sliding window analysis. Progeny along with an uncloned sample (denoted with an *) are shown as rows and each column represents a 90kb region (window size was defined as the expected distance between heterozygous SNP calls based on the heterozygous SNP call rate for each cross). (A) In progeny of the NF54 x NHP4026 cross, 25 progeny had regions with above expected heterozygosity. (B) In progeny of the MKK2835 x NHP1337 cross, 35 progeny had regions with above expected heterozygosity. The un-cloned samples (*) show above background heterozygosity or high heterozygosity across the genome.

**S1 Table. Mosquito stage crossing results.**

**S2 Table. Cloning methodology and results.**

**S3 Table. NF54/NF54-GFPLuc x NHP4026 Genetic Map**

**S4 Table. MKK2835 x NHP1337 Genetic Map**

**S5 Table. Allele frequencies and significance of segregation distortion in NF54 x NHP4026 progeny.**

**S6 Table. Microsatelite information.**

## References

1. Fairlamb AH, Gow NAR, Matthews KR, Waters AP. Drug resistance in eukaryotic microorganisms. Nature Microbiology. 2016;1(7). doi: 10.1038/nmicrobiol.2016.92.

2. Wang W, Wang L, Liang Y-S. Susceptibility or resistance of praziquantel in human schistosomiasis: a review. Parasitology Research. 2012;111(5):1871–7. doi: 10.1007/s00436-012-3151-z.

3. Ferdig MT, Cooper RA, Mu J, Deng B, Joy DA, Su XZ, et al. Dissecting the loci of low-level quinine resistance in malaria parasites. Mol Microbiol. 2004;52(4):985–97. doi: 10.1111/j.1365-2958.2004.04035.x“.

4. Müller IB, Hyde JE. Antimalarial drugs: modes of action and mechanisms of parasite resistance. Future Microbiology. 2010;5(12):1857–73. doi: 10.2217/fmb.10.136.

5. Chevalier FD, Valentim CLL, LoVerde PT, Anderson TJC. Efficient linkage mapping using exome capture and extreme QTL in schistosome parasites. BMC Genomics. 2014;15(1). doi: 10.1186/1471-2164-15-617.

6. Alsford SAM, Kelly JM, Baker N, Horn D. Genetic dissection of drug resistance in trypanosomes. Parasitology. 2013;140(12):1478–91. doi: 10.1017/s003118201300022x.

7. Miotto O, Amato R, Ashley EA, MacInnis B, Almagro-Garcia J, Amaratunga C, et al. Genetic architecture of artemisinin-resistant Plasmodium falciparum. Nature genetics. 2015;47(3):226–34.

8. Ariey F, Witkowski B, Amaratunga C, Beghain J, Langlois A-C, Khim N, et al. A molecular marker of artemisinin-resistant Plasmodium falciparum malaria. Nature. 2014;505(7481):50–5.

9. Straimer J, Gnadig NF, Witkowski B, Amaratunga C, Duru V, Ramadani AP, et al. Drug resistance. K13-propeller mutations confer artemisinin resistance in Plasmodium falciparum clinical isolates. Science (New York, NY). 2015;347(6220):428–31. doi: 10.1126/science.1260867 [doi].

10. Cheeseman IH, Miller BA, Nair S, Nkhoma S, Tan A, Tan JC, et al. A major genome region underlying artemisinin resistance in malaria. Science (New York, NY). 2012;336(6077):79–82. doi: 10.1126/science.1215966 [doi].

11. Fidock DA, Nomura T, Talley AK, Cooper RA, Dzekunov SM, Ferdig MT, et al. Mutations in the P. falciparum digestive vacuole transmembrane protein PfCRT and evidence for their role in chloroquine resistance. Mol Cell. 2000;6(4):861–71. Epub 2000/11/25. PubMed PMID: 11090624; PubMed Central PMCID: PMCPMC2944663.

12. Valentim CLL, Cioli D, Chevalier FD, Cao X, Taylor AB, Holloway SP, et al. Genetic and Molecular Basis of Drug Resistance and Species-Specific Drug Action in Schistosome Parasites. Science. 2013;342(6164):1385–9. doi: 10.1126/science.1243106.

13. Sá JM, Kaslow SR, Krause MA, Melendez-Muniz VA, Salzman RE, Kite WA, et al. Artemisinin resistance phenotypes and K13 inheritance in a Plasmodium falciparum cross and Aotus model. Proceedings of the National Academy of Sciences. 2018. doi: 10.1073/pnas.1813386115.

14. MacLeod A. The genetic map and comparative analysis with the physical map of Trypanosoma brucei. Nucleic Acids Research. 2005;33(21):6688–93. doi: 10.1093/nar/gki980.

15. Cheeseman IH, McDew-White M, Phyo AP, Sriprawat K, Nosten F, Anderson TJ. Pooled sequencing and rare variant association tests for identifying the determinants of emerging drug resistance in malaria parasites. Mol Biol Evol. 2015;32(4):1080–90. Epub 2014/12/24. doi: 10.1093/molbev/msu397. PubMed PMID: 25534029; PubMed Central PMCID: PMCPMC4379400.

16. Su X, Ferdig MT, Huang Y, Huynh CQ, Liu A, You J, et al. A genetic map and recombination parameters of the human malaria parasite Plasmodium falciparum. Science. 1999;286(5443):1351–3. Epub 1999/11/13. PubMed PMID: 10558988.

17. Jiang H, Li N, Gopalan V, Zilversmit MM, Varma S, Nagarajan V, et al. High recombination rates and hotspots in a Plasmodium falciparum genetic cross. Genome Biol. 2011;12(4):R33. Epub 2011/04/06. doi: 10.1186/gb-2011-12-4-r33. PubMed PMID: 21463505; PubMed Central PMCID: PMCPMC3218859.

18. Gardner MJ, Hall N, Fung E, White O, Berriman M, Hyman RW, et al. Genome sequence of the human malaria parasite Plasmodium falciparum. Nature. 2002;419(6906):498–511. doi: 10.1038/nature01097. PubMed PMID: 12368864; PubMed Central PMCID: PMCPMC3836256.

19. Aurrecoechea C, Brestelli J, Brunk BP, Dommer J, Fischer S, Gajria B, et al. PlasmoDB: a functional genomic database for malaria parasites. Nucleic acids research. 2009;37(Database issue):D539–43. doi: 10.1093/nar/gkn814 [doi].

20. Aurrecoechea C, Barreto A, Basenko EY, Brestelli J, Brunk BP, Cade S, et al. EuPathDB: the eukaryotic pathogen genomics database resource. Nucleic Acids Research. 2017;45(D1):D581–D91. doi: 10.1093/nar/gkw1105.

21. Miles A, Iqbal Z, Vauterin P, Pearson R, Campino S, Theron M, et al. Indels, structural variation, and recombination drive genomic diversity in Plasmodium falciparum. Genome Res. 2016;26(9):1288–99. doi: 10.1101/gr.203711.115. PubMed PMID: 27531718; PubMed Central PMCID: PMCPMC5052046.

22. Flint J, Mackay TFC. Genetic architecture of quantitative traits in mice, flies, and humans. Genome Research. 2009;19(5):723–33. doi: 10.1101/gr.086660.108.

23. Vaughan AM, Pinapati RS, Cheeseman IH, Camargo N, Fishbaugher M, Checkley LA, et al. Plasmodium falciparum genetic crosses in a humanized mouse model. Nature Methods. 2015;12(7):631–3. doi: 10.1038/nmeth.3432.

24. Mott BT, Eastman RT, Guha R, Sherlach KS, Siriwardana A, Shinn P, et al. High-throughput matrix screening identifies synergistic and antagonistic antimalarial drug combinations. Scientific reports. 2015;5:13891. doi: 10.1038/srep13891 [doi].

25. Hayton K, Gaur D, Liu A, Takahashi J, Henschen B, Singh S, et al. Erythrocyte Binding Protein PfRH5 Polymorphisms Determine Species-Specific Pathways of Plasmodium falciparum Invasion. Cell Host & Microbe. 2008;4(1):40–51. doi: 10.1016/j.chom.2008.06.001.

26. Logan-Klumpler FJ, De Silva N, Boehme U, Rogers MB, Velarde G, McQuillan JA, et al. GeneDB--an annotation database for pathogens. Nucleic Acids Res. 2012;40(Database issue):D98–108. Epub 2011/11/26. doi: 10.1093/nar/gkr1032. PubMed PMID: 22116062; PubMed Central PMCID: PMCPMC3245030.

27. Okamoto N, Spurck TP, Goodman CD, McFadden GI. Apicoplast and Mitochondrion in Gametocytogenesis ofPlasmodium falciparum. Eukaryotic Cell. 2009;8(1):128–32. doi: 10.1128/ec.00267-08.

28. Vaidya AB, Morrisey J, Plowe CV, Kaslow DC, Wellems TE. Unidirectional dominance of cytoplasmic inheritance in two genetic crosses of Plasmodium falciparum. Molecular and Cellular Biology. 1993;13(12):7349–57. doi: 10.1128/mcb.13.12.7349.

29. Creasey AM, Ranford-Cartwright LC, Moore DJ, Williamson DH, Wilson RJ, Walliker D, et al. Uniparental inheritance of the mitochondrial gene cytochrome b in Plasmodium falciparum. Curr Genet. 1993;23(4):360–4. Epub 1993/01/01. doi: 10.1007/bf00310900. PubMed PMID: 8467535.

30. Ranford-Cartwright LC, Mwangi JM. Analysis of malaria parasite phenotypes using experimental genetic crosses of Plasmodium falciparum. International Journal for Parasitology. 2012;42(6):529–34. doi: 10.1016/j.ijpara.2012.03.004. PubMed PMID: WOS:000305720200003.

31. Gabryszewski SJ, Modchang C, Musset L, Chookajorn T, Fidock DA. Combinatorial Genetic Modeling ofpfcrt-Mediated Drug Resistance Evolution inPlasmodium falciparum. Molecular Biology and Evolution. 2016;33(6):1554–70. doi: 10.1093/molbev/msw037.

32. Veiga MI, Osorio NS, Ferreira PE, Franzen O, Dahlstrom S, Lum JK, et al. Complex polymorphisms in the Plasmodium falciparum multidrug resistance protein 2 gene and its contribution to antimalarial response. Antimicrob Agents Chemother. 2014;58(12):7390–7. Epub 2014/10/01. doi: 10.1128/AAC.03337-14. PubMed PMID: 25267670; PubMed Central PMCID: PMCPMC4249497.

33. Li X, Kumar S, McDew-White M, Haile M, Cheeseman IH, Emrich S, et al. Genetic mapping of fitness determinants across the malaria parasite Plasmodium falciparum life cycle. PLoS Genet. 2019;15(10):e1008453. Epub 2019/10/15. doi: 10.1371/journal.pgen.1008453. PubMed PMID: 31609965; PubMed Central PMCID: PMCPMC477818.

34. Parija S, Antony H. Antimalarial drug resistance: An overview. Tropical Parasitology. 2016;6(1). doi: 10.4103/2229-5070.175081.

35. Nzila A, Mwai L. In vitro selection of Plasmodium falciparum drug-resistant parasite lines. J Antimicrob Chemother. 2010;65(3):390–8. Epub 2009/12/22. doi: 10.1093/jac/dkp449. PubMed PMID: 20022938; PubMed Central PMCID: PMCPMC2818104.

36. Morgante F, Huang W, Maltecca C, Mackay TFC. Effect of genetic architecture on the prediction accuracy of quantitative traits in samples of unrelated individuals. Heredity. 2018;120(6):500–14. doi: 10.1038/s41437-017-0043-0.

37. Michel K, Morlais I, Nsango SE, Toussile W, Abate L, Annan Z, et al. Plasmodium falciparum Mating Patterns and Mosquito Infectivity of Natural Isolates of Gametocytes. Plos One. 2015;10(4). doi: 10.1371/journal.pone.0123777.

38. Ranford-Cartwright LC. Fit for fertilization: Mating in malaria parasites. Parasitology Today. 1995;11(4):154–7. doi: 10.1016/0169-4758(95)80138-3.

39. Ranford-Cartwright LC, Balfe P, Carter R, Walliker D. Frequency of cross-fertilization in the human malaria parasite Plasmodium falciparum. Parasitology. 2009;107(01). doi: 10.1017/s003118200007935x.

40. Walliker D, Quakyi IA, Wellems TE, McCutchan TF, Szarfman A, London WT, et al. Genetic analysis of the human malaria parasite Plasmodium falciparum. Science. 1987;236(4809):1661–6. Epub 1987/06/26. PubMed PMID: 3299700.

41. Walker-Jonah A, Dolan SA, Gwadz RW, Panton LJ, Wellems TE. An RFLP map of the Plasmodium falciparum genome, recombination rates and favored linkage groups in a genetic cross. Molecular and Biochemical Parasitology. 1992;51(2):313–20. doi: 10.1016/0166-6851(92)90081-t.

42. Myburg AA, Vogl C, Griffin AR, Sederoff RR, Whetten RW. Genetics of postzygotic isolation in Eucalyptus: whole-genome analysis of barriers to introgression in a wide interspecific cross of Eucalyptus grandis and E. globulus. Genetics. 2004;166(3):1405–18. Epub 2004/04/15. PubMed PMID: 15082559; PubMed Central PMCID: PMCPMC1470765.

43. Yin TM, DiFazio SP, Gunter LE, Riemenschneider D, Tuskan GA. Large-scale heterospecific segregation distortion in Populus revealed by a dense genetic map. Theor Appl Genet. 2004;109(3):451–63. Epub 2004/05/29. doi: 10.1007/s00122-004-1653-5. PubMed PMID: 15168022.

44. Bodenes C, Chancerel E, Ehrenmann F, Kremer A, Plomion C. High-density linkage mapping and distribution of segregation distortion regions in the oak genome. DNA Res. 2016;23(2):115–24. Epub 2016/03/26. doi: 10.1093/dnares/dsw001. PubMed PMID: 27013549; PubMed Central PMCID: PMCPMC4833419.

45. Tirrell AR, Vendrely KM, Checkley LA, Davis SZ, McDew-White M, Cheeseman IH, et al. Pairwise growth competitions identify relative fitness relationships among artemisinin resistant Plasmodium falciparum field isolates. Malar J. 2019;18(1):295. Epub 2019/08/30. doi: 10.1186/s12936-019-2934-4. PubMed PMID: 31462253; PubMed Central PMCID: PMCPMC6714446.

46. Anthony TG, Polley SD, Vogler AP, Conway DJ. Evidence of non-neutral polymorphism in Plasmodium falciparum gamete surface protein genes Pfs47 and Pfs48/45. Mol Biochem Parasitol. 2007;156(2):117–23. Epub 2007/09/11. doi: 10.1016/j.molbiopara.2007.07.008. PubMed PMID: 17826852.

47. Manske M, Miotto O, Campino S, Auburn S, Almagro Garcia J. Analysis of Plasmodium falciparum diversity in natural infections by deep sequencing. Nature. 2012;487(7407):375–9. doi: 10.1038/nature11174.

48. Xu S, Hu Z. Mapping Quantitative Trait Loci Using Distorted Markers. International Journal of Plant Genomics. 2009;2009:1–11. doi: 10.1155/2009/410825.

49. Li HW. Aligning sequence reads, clone sequences and assembly contigs with BWA-MEM. arXiv:13033997v2. 2013.

50. DePristo MA, Banks E, Poplin R, Garimella KV, Maguire JR, Hartl C, et al. A framework for variation discovery and genotyping using next-generation DNA sequencing data. Nature Genetics. 2011;43(5):491–8. doi: 10.1038/ng.806.

