## Supplementary material for "Surprising variation in the outcome of two malaria genetic crosses using humanized mice: implications for genetic mapping and malaria biology": S1 File

**Cloning by limiting dilution: plate set up, screening and stock/sample generation**

November 29, 2017

Lisa Checkley Needham

*Having fresh blood (less than a few days from donation) is critical to cloning success. Plan ahead to have fresh blood delivered right before cloning.

1) Cloning plate set up:

Preparation of Initial Culture

Thawed parasites: Thaw parasite line/stock as per lab protocol. Depending on the needs of the project can either:

-thaw in the morning and check late in the day/early next morning and set up

plates based on parasitemia (this would work for any straight from mouse blood)

-put on shaker right away and let go a minimum of 2 life cycles (to reduce number	of multiple invasions) and then make a slide for parasitemia

*sometimes lines that are not good growers require a life cycle or two without

shaking to get established before putting on the shaker.

After two life cycles on shaker (or if it is direct from mouse blood skip to this step) make a smear and count parasitemia.

Straight from mouse: If parasitemia is 0.25 or higher set-up a few cloning plates immediately after *in vitro* transition. Otherwise allow cultures to establish in small volume cultures on the shaker and set-up cloning plates as soon as parasites are growing well, but as cultures get established make sure to expand the culture as soon as parasitemia reaches 1%. Do not let cultures reinvade at or above 1%.

Cloning Plate dilutions

Set up plates as follows for 0.25 parasites/well:

**Dilution 1**=%Parasitemia/2 (this is the volume of complete media in mL you add to a tube) + 10uL of culture. Mix well. For example: for a culture at 1% parasitemia, add 500 uL of CM to 10 uL of culture.

**Dilution 2**=10 uL of Dilution 1 + 1 mL of complete media in a tube and mix well

**Diluiton3**=Add 200 uL of Dilution 2 to 38mLs of complete media + 2 mLs of RBC suspension (we use 50/50 rbc and incomplete media, use fresh RBCs). Mix well.

Plate 200 uL of the above dilution into the wells of 96 well plate with a mulit-channel pipette using a new set of tips for each row. Put the plates into a clean chamber with a 96 well plate of H_2_O in the bottom for humidity.

To maximize returns we can use a larger amount of original culture and go as high as 0.5 parasites/well. At this level we have observed 10-20% of positive wells are non-clonal. We can then sub-clone to get additional parasites.

Be sure to start cloning plates in as fresh of blood as possible (maximum of a few days old)

Cloning Plate Maintenance

Change media on plates weekly by removing 150 uL of media (use new tips for each row-one box of tips per plate) and replace with 150 uLs of a mixture of 30 mLs complete media + 400 uLs of RBCs that has been mixed well. As plates age if media begins to look dark and RBC lysis increases, change the media more frequently (every 4-5 days) to ensure parasites have a fresh supply of RBCs.

Gas the chambers every 2 days.

While changing media on plates, clean the chamber out with 10%	bleach water and also clean the plate with the water in the bottom of the chamber. I refill the plate for humidity and put it and the clean disassembled chamber into a hood and UV it for 30 mins. I also use pipettors that are dedicated to cloning use only to minimize contamination

2) Screening cloning plates:

kit for PCR: **Phusion Blood Direct PCR Kit Thermo Scientific cat# F-547L**. I recommend taking the time to go to their primer calculator to see what temp to use with this kit. It is rare that I get the same TM to use for my primers with this kit as what is recommended by IDT (where we order most of our primers). People in the lab who have ignored this have not had good luck with this kit.

**Invitrogen/Life Sciences/Molecular Probes SYBR Green I nucleic acid stain 10,000X cat # S7585**. Dilute this in water to ~3X concentration (since once it goes into reaction will be ~1X in a tube and use that as water for reactions).

**Primers** – Can use any primers with a small amplicon (<100 bp) that are well behaved. We recommend either

-the microsatellite primer PE14D which means our annealing temp is 53°, we use 54°for the qPCR and it works fine)

-pfCRT primers

**Master Mix**

-Make a master mix for plates. We use an Eppendorf robot to move it into test plates. Make up about an extra plate’s worth of master mix in order to use the robot. The cost of the reagents is minimal next to the time it saves. Use the following for 4 test plates:

1RXN:	500 RXN Master mix:

2X buffer	5.0 uL	2500 uL

enzyme	0.1 uL	50 uL

F. Primer	0.25 uL	125 uL

R. Primer	0.25 uL	125 uL

H_2_O (w/SYBR)	1.4 uL	700 uL

7.0 uL	3500 uL

**Test Plate**

-add 7 uL of the master mix to the test plate

-dilute whole culture 1:4 (1 uL of whole culture from cloning plates into 3 uL of water) and add 3 uL of diluted to the test plate (I do this right after I have changed the media so the wells are already mixed up).

-seal the plate and run on the qPCR protocol

**PCR Protocol**

- ABI 7900HT for pfCRT primers

stage 1=95° for 20 seconds

stage 2=95° for 1 second

30 cycles

62.3° for 30 seconds

65° for 15 seconds

-set to “fast” mode, reaction volume= 10 uL, and “SYBR detector no quencher”

-plate run time is between 30-40 mins (my last run took 33).

-after the run and analysis use the CT score to determine positives (though usually pretty obvious by looking at the curves. See images below). Doug will sort based on score and then look at the CT numbers. We have found that CT=21 or less is a decent cutoff. Some at 21 will be true positives and some will not. Anything with a CT=21 or lower will look at the individual curve and decide. Because the kit is so sensitive it is possible to leave questionable wells and screen them again the next week. Usually one more week will make it more obvious if it is a true positive.

A lot depends on the threshold set by the software automatically and can vary from plate to plate.

*curves in oval are positive wells (except one greenish one)

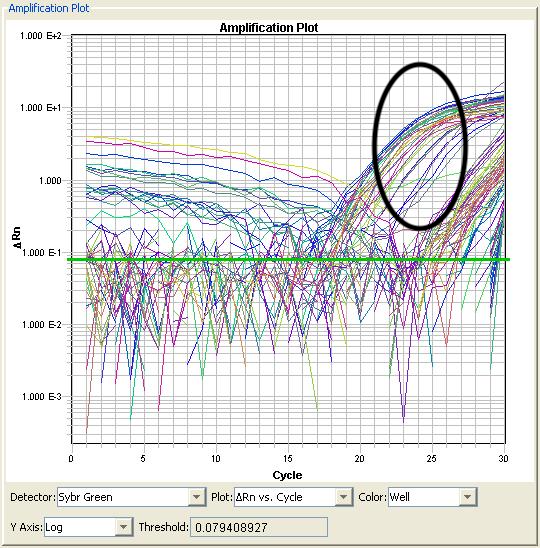

*Two curves on the left are positives, two on the right are negatives. Less busy plot.

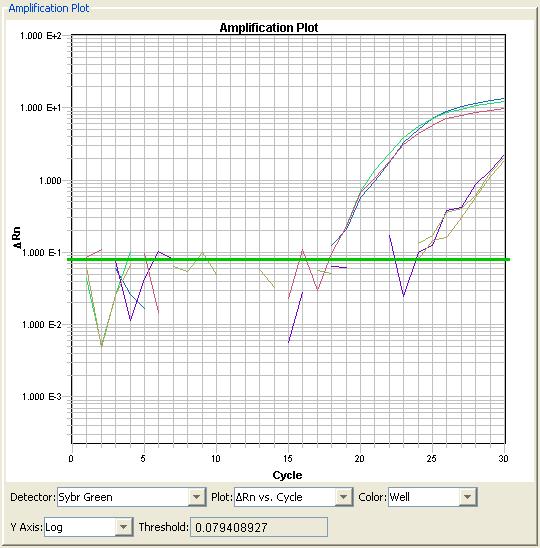

One last example on the next page: both these curves have about the same CT but only the one on the left is positive. So we don’t rely solely on CT, we look at the curves of all potential positives and then decide which ones we think are real. If something seems questionable we leave it till the next screen.

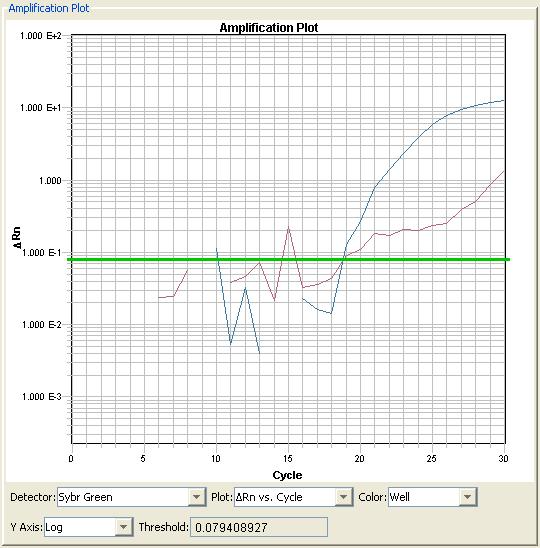

Positives are pulled to 1 mL wells at 5% Hct in 24 well plates and maintained at standard culture conditions to generate stock pellets and pellets for DNA isolation.

Stock Pellets: Parasites need to be greater than 1% parasitemia and mostly ring.

DNA Pellets: Let the wells grow up until the parasitemia is above 3% and late stage then freeze the whole well/culture down for DNA.
